# Global delimitation of *Cyanoboletus, Cacaoporus* and *Cupreoboletus* (Suillelloideae : Boletaceae)

**DOI:** 10.64898/2026.05.12.724631

**Authors:** Paulo Oliveira, Rodrigo Mariquito

**Author notes:** Corresponding author: Paulo Oliveira.

## Abstract

This investigation aimed at compiling all phylogenetic lineages within and around genus *Cyanoboletus*. The evolutionary inference obtained from the nuclear rDNA internal transcribed spacer region (ITS) suggests excluding part of the species currently classified in *Cyanoboletus*, leaving only the species that develop a strong staining reaction to touch and to air exposure of the context. The excluded lineages are the monotypic *Cupreoboletus* genus and a few species that do not develop such reaction, the latter forming a clade together with genera *Cacaoporus* and *Acyanoboletus*, thus broadening the concept of *Cacaoporus* to encompass all of them. The emerging ‘3C’ perspective of *Cupreoboletus, Cacaoporus* and *Cyanoboletus* offers a remarkably consistent morphological diagnosis, overcoming the problems of a too broad concept for *Cyanoboletus. Boletus neotropicus, B. novae-zelandiae* and *B. sensibilis* belong respectively in *Cyanoboletus, Cacaoporus* and *Lanmaoa*, and lineages are indicated that probably represent undescribed species, at least four in *Cacaoporus* and at least five in *Cyanoboletus*. Diagnostic tables and dichotomic keys are presented by geographic region. The present work also includes a study of the phylogenetic position of *Neoboletus flavosanguineus*, a species once classified in *Cyanoboletus*. The complexity of assigning species epithets in some lineages is addressed, namely for the boundaries between *Cacaoporus instabilis* and *Ca. fagaceophilus* as well as the diversity under the names *Cyanoboletus sinopulverulentus* and *Cy. pulverulentus*. The overall picture of evolutionary lineages sets a framework for the choice of reference data that can provide a balanced and efficient coverage in future phylogenetic studies that involve the 3C.

## Introduction

*Cyanoboletus* Gelardi, Vizzini & Simonini is a genus in family Boletaceae Chevall. erected in 2014 around *Boletus pulverulentus* Opat., the type species of *Boletus* sect. *Subpruinosi* Fr., as part of the progressive clarification of the phylogenetic relationships within the ‘classical’ concept of *Boletus* L. (Binder and Hibbett 2006, Wu et al. 2014, Magnago et al. 2022; Tremble et al. 2024). Currently, the prevailing understanding places *Cyanoboletus* in the subfamily Suillelloideae Dentinger, Tremble, Halling, T.W. Henkel & Moncalvo (Tremble et al. 2023) containing all ‘classical’ sections of *Boletus* except for the current *Boletus sensu stricto* (formerly *Boletus* sect. *Edules* Fr.), together with many other genera, some containing long-known European species (*Baorangia* G. Wu & Zhu L. Yang, *Butyriboletus* D. Arora & J.L. Frank, *Caloboletus* Vizzini, *Cupreoboletus* Simonini, Gelardi & Vizzini, *Exsudoporus* Vizzini, Simonini & Gelardi, *Imperator* Koller, Assyov, Bellanger, Bertéa, Loizides, G. Marques, P.-A. Moreau, J.A. Muñoz, Oppicelli, D. Puddu & F. Rich., *Lanmaoa* G. Wu & Zhu L. Yang, *Neoboletus* Gelardi, Simonini & Vizzini, *Pulveroboletus* Murrill, *Rubroboletus* Kuan Zhao & Zhu L. Yang and *Suillellus* Murrill), joined by a plethora of genera known only outside Europe (*Acyanoboletus* G. Wu & Zhu L. Yang, *Amoenoboletus* G. Wu, E. Horak & Zhu L. Yang, *Cacaoporus* Raspé & Vadthanarat, *Costatisporus* T.W. Henkel & M.E. Sm., *Crocinoboletus* N.K. Zeng, Zhu L. Yang & G. Wu, *Erythrophylloporus* Ming Zhang & T.H. Li, *Gastroboletus* Lohwag, *Gymnogaster* J.W. Cribb, *Hongoboletus* G. Wu & Zhu L. Yang, *Pseudobaorangia* D.F. Sun, R. Hua, F. Zhou & J.B. Zhang, *Rugiboletus* G. Wu & Zhu L. Yang, *Singerocomus* T.W. Henkel & M.E. Sm., *Sutorius* Halling, Nuhn & N.A. Fechner and *Tropicoboletus* Angelini, Gelardi & Vizzini). Some authors criticise this proliferation of genera names, and merging of genera has already been proposed — but not without countercriticism — therefore the taxonomic arrangements in the Suillelloideae are expected to evolve, as new insights are gained (Wu et al. 2016; Wang et al. 2024; Biketova et al. 2026).

Currently 17 taxa at species level have been classified in *Cyanoboletus*, forming a relatively small but expanding assembly (Table 1).

**Table 1.**
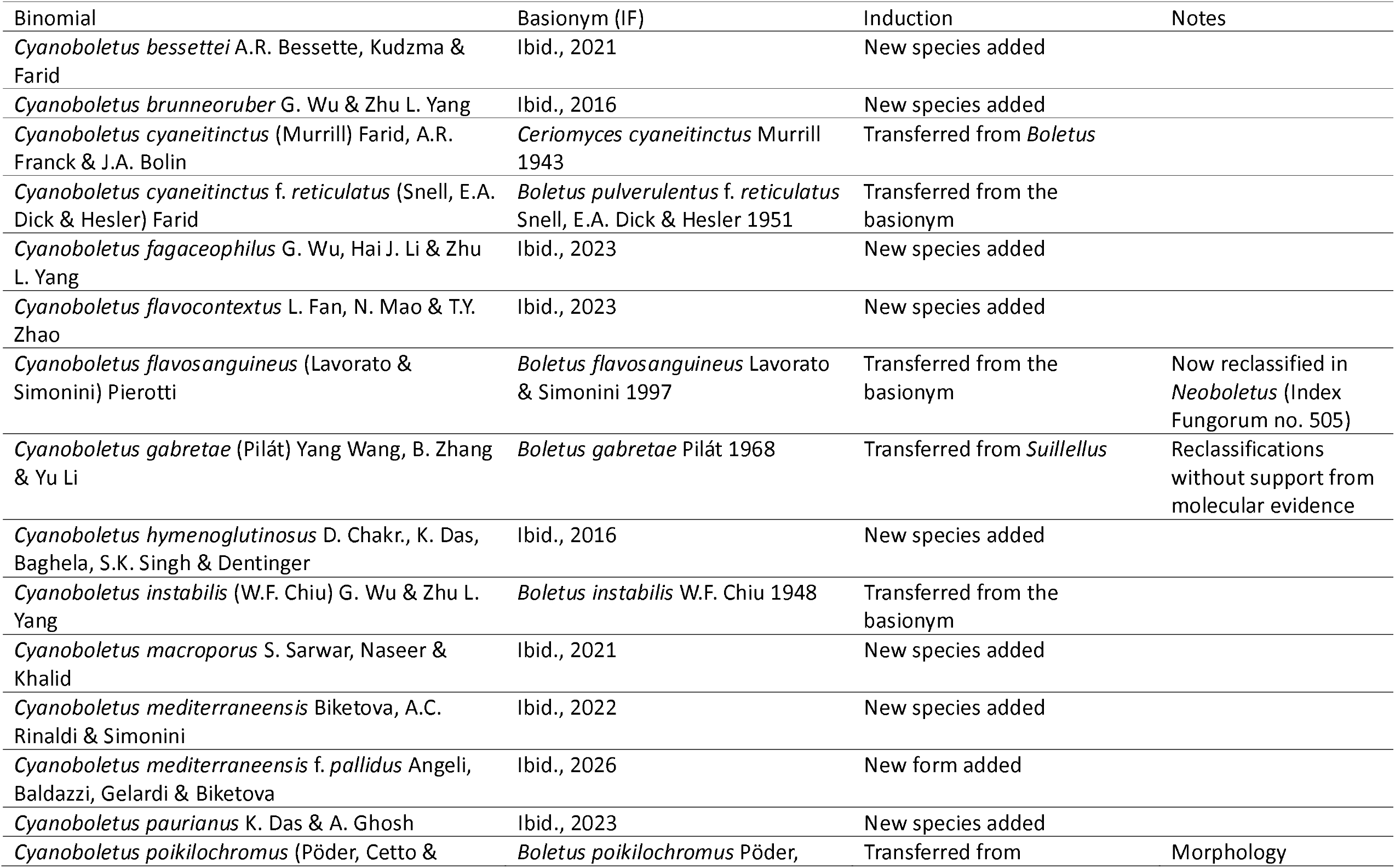
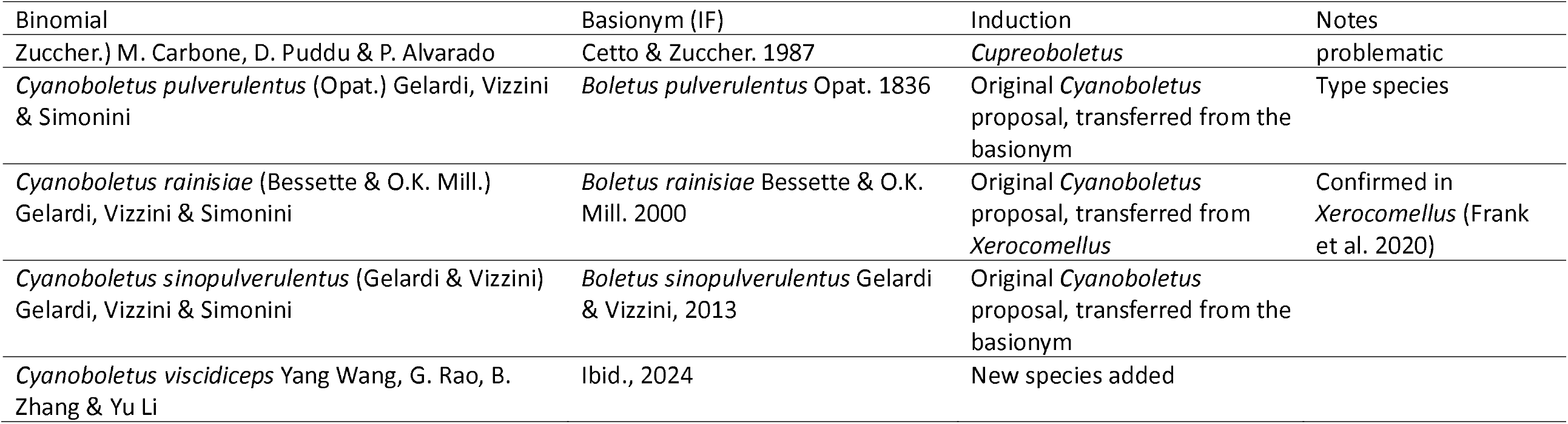
*Cyanoboletus* species according to *Index Fungorum* (May 12, 2026).

The *Cyanoboletus* name relates to the type species showing a prompt dark blue staining of any part of the surface by touch or bruising, and of the context by exposure to air (Vizzini 2014), but this character is not in itself defining, as it is prevalent in genera such as *Imperator* and *Neoboletus* (Muñoz 2005). Additional diagnostic characters include the brownish pileus at maturity, the relatively small ‘xerocomoid’ habit with a non-ventricose stipe lacking developed reticulum (if present, incipient as if a short prolongation of the hymenophore), smooth spores and hymenophoral trama of the boletoid type (Wu et al. 2016).

The inclusion of *Boletus poikilochromus* in *Cyanoboletus* (Carbone et al. 2023) is but one example where the results of multigene phylogenies shatter established morphological concepts. Such phylogenies are based on genomic regions that do not contribute to a stable dendrogram topology by themselves, but often combine their ‘phylogenetic signal’ such that improved resolution and clade support is achieved (Edwards et al. 2007); thus, the multigene approach is well-established in mycological taxonomy and, in general, is deemed good enough to resolve broad intergeneric relationships, even if the chosen genes vary from one study to another. A very recent example of this practice could not fail to repeat the placement of that species in *Cyanoboletus* (Biketova et al. 2026).

On the other hand, the nuclear rDNA internal transcribed spacers (ITS), a crucial ‘barcode’ for fungi (Xu 2016), tends to be relegated in such studies, which is understandable because of its fast evolutionary divergence marked by extensive insertions and deletions that pose major difficulties with aligning nucleotide sequences except for relatively narrow taxonomic scopes. Unfortunately, many taxonomic descriptions of new species of fungi forgo the sequencing of this barcode and its submission to public databases, missing the opportunity to support metabarcoding research or the increasing use of BLAST searches by citizen science projects.

Protein-coding genes are selectively constrained, sometimes also in their introns (Nei and Kumar 2000), and the selection-neutral character of the ITS offers a cross-validation of the taxonomic results of every multigene analysis and should be worthwhile to be attempted (Frank et al. 2020; Oliveira and Arraiano-Castilho 2025). The present work addresses phylogenetic relationships, based on a comprehensive sampling of DNA sequences available in public databases, that pertain to *Cyanoboletus pulverulentus* and its allies. A novel intergeneric boundary is presented — featuring a significant expansion of genus *Cacaoporus* to include some of the species listed in Table 1 — and its implication for morphological diagnosis is realized in regional diagnoses for known species.

## Material & Methods

### Compilation of sequences

An initial search was made in the NCBI Taxonomy page https://www.ncbi.nlm.nih.gov/datasets/taxonomy/browser/ (O’Leary et al. 2024) to build the dataset for the Suillelloideae (Online Resource 1). The genes included in the searches were the ITS, the nuclear ribosomal DNA large/28S subunit (nucLSU), the two largest subunits of RNA polymerase II (RPB1 and RPB2) and the translation elongation factor 1 α subunit (TEF1), as they are the ones providing the greatest species coverage in this subfamily. Subsequently, a selection of ITS sequences classified as *Cyanoboletus* in the obtained Suillelloideae phylogeny (Online Resource 2) were used in a MassBlaster search in the PlutoF workbench (Abarenkov et al. 2010) to sample sequences by their proximity to the named ones, including from environmental samples. A similar sampling was made with non-ITS sequences via the NCBI Nucleotide BLAST portal (blastn suite, https://blast.ncbi.nlm.nih.gov/). A search in the European Nucleotide Archive (https://www.ebi.ac.uk/ena/browser/sequence-search) yielded one sequence apparently not represented in Genbank (accession KIGB01028760). Except for the sequences of *Acyanoboletus rugosipileus*, released later (on our request, April 19, 2026), and for the extra sequences retrieved for the study focused on *Neoboletus flavosanguineus* (April 24, 2026), all searches ended on March 18, 2026 (Online Resource 1).

### Alignments and data sessions

All alignments except for the ITS dataset were made in UGene (Okonechnikov et al. 2012), using the MUSCLE tool version 5 (Edgar 2022), supplemented by visual inspections and corrections of obvious misalignments in the automated result.

Aligned datasets were concatenated in MEGA12 (Kumar et al. 2024) and saved as FastA files, as well as the corresponding ‘data session’ files for future examination. All sequences in single-gene phylogenies are identified by their accession number and title as used in Genbank at the time of their retrieval.

In the case of the ITS dataset, a small group of sequences from the different genera were selected from the automatic alignment with MUSCLE and submitted to the ITSx tool (Nilsson et al. 2010) in the PlutoF workbench to determine the limits for the 5.8S gene, such that the already existing ITS alignment was partitioned into the ITS1, 5.8S and ITS2 regions, and each partition exported without gaps (that is, reversing to the aligned sequences) to separate FastA files so that they could be treated as different partitions. Prior to using them in the bali-phy application version 4.1 (Redelings 2021) the following accessions were excluded for having short sequences and being almost redundant to others that were satisfactory: MG030473, MW376644, MW675732, MW675733 and ON705304 (they are nevertheless listed and classified in Online Resource 1). Bali-phy offers a combined Bayesian estimation of multiple sequence alignments and phylogenetic trees, outperforming other automatic alignment tools especially when evolutionary distances are larger (Redelings 2014). The command used (WSL Ubuntu environment) was «bali-phy -c ITSops.txt -S ‘tn93 +> Rates.gamma +> inv’ &», with settings (in ITSops.txt) specifying the input files and allowing a separate modelling of the three partitions, but linking the intronic sequences ITS1 and ITS2. The tn93 parameter stands for the Tamura-Nei substitution model (Tamura and Nei 1993). After initial shorter runs to evaluate the evolution of the Bayesian inference, a final run was conducted, lasting 455.46 hours in a laptop with an Intel Core i5-8365U CPU @ 1.60GHz (1.90 GHz), with 8 GB Random Access Memory. Using the bp-analyze command, an MCMC Post-hoc Analysis was generated, reporting 17930 samples, a burnin of 1992, and ESS of 311 for likelihood (with 498 burnin). Of all the statstics reported, only |A| (number of alignment columns) was at a less than satisfactory level (ESS 81 for the ITS2, ESS 63 overall). The best (WPD) alignments for the three partitions were concatenated with the command alignment-cat to generate the final alignment. The greedy consensus tree with posterior probability annotations was retrieved as output tree to be processed as the remainder trees (see below).

### Phylogenetic analyses

#### Preliminary work

Quick evaluations of the MUSCLE alignments were made by building phylogenetic splits networks in Splitstree version 6.4 through 6.7 (Huson and Bryant 2006). Tanglegrams were built in Dendroscope (Huson and Scornavacca 2012) to compare alternative versions of a phylogenetic tree, especially when the performance of different phylogenetic models was under inspection. Based on this procedure, it was determined that for BEAST (detailed below) it was preferable to use a strict clock with single-gene alignments, and a relaxed clock with concatenate alignments.

#### Pairwise reconstructions and statistics

The sequences in the ITS alignment were grouped by putative species in MEGA12, and this framework used for two sets of analyses. First, a phylogenetic reconstruction using the Minimum Evolution method (Rhetsky and Nei 1992), with uncorrected p-distances and pairwise deletion, for evaluation of the interspecies gap, by collapsing the branches in the phylogenetic reconstruction according to a variety of similarity thresholds in the resulting ‘tree session’ (Collapse Nodes > by Sequence Difference). Second, for estimations of the mean within- and between-group p-distances (Nei and Kumar 2000).

#### Bayesian phylogenetic reconstructions

Except for the ITS dataset, where the tree was obtained as part of the alignment process (see above), the BEAST package version 10.5.0 (Baele et al. 2025b) was used as follows: an alignment was introduced as a single partition in the BEAUti tool, modelling the analysis to be made with the nucleotide substitution model GTR+gamma (equal weights)+invariant sites with 5 gamma categories (Tavaré 1986, Yang 1994), strict clock or the Hamiltonian Monte Carlo relaxed clock with Lognormal distribution (Baele et al. 2025a), a Speciation Birth-Death process tree prior (Gernhard 2008; Magee et al. 2020), the UPGMA starting tree, and a MCMC 10^7^ chain length, with parameter logging every 10^3^ states. Individual variations of this model are detailed in the legend for each tree. The XML file produced by BEAUti was then analysed with BEAST, supported by the BEAGLE library version 4 (Ayres et al. 2019). Two BEAST runs were always performed in parallel. Using the external Tracer tool version 1.7.2 (Rambaut et al. 2018), the convergence of each and both runs was inspected, and the best run according to the ESS joint parameter was selected for further processing in the Treeannotator tool. The latter used the MrHIPSTR — Majority rule highest independent posterior subtree reconstruction (Baele et al. 2025a) with a burnin of at least 10^6^ states and Median node heights. Individual variations of this procedure are detailed in the legend for each tree. In the case of the ITS+nucLSU phylogeny, the BEAST output trees (2 x 10^7^ states for each run) were combined using Logcombiner with a 2 x 10^6^ burnin each, and the resulting set was processed in Treeannotator without burnin.

Each final tree selected by Treeannotator was uploaded to iTOL version 7 (Letunic and Bork 2024) for final rendering, rooting in *Pulchroboletus* (Gelardi et al. 2023). For clarity, all posterior probability support values below 70% are not shown.

The bali-phy output, all alignment FastA files, MEGA12 data session files, BEAUti XML files, and NEXUS tree files are available in the Figshare link doi: 10.6084/m9.figshare.32298045 (Oliveira and Mariquito 2026).

## Results

Our own multigene phylogeny of the Suillelloideae has confirmed previous results (clades 49-50 in Wu et al. 2016; Gelardi et al. 2023; Tremble et al. 2024) suggesting a subgroup, with 88% posterior probability, encompassing the genera *Lanmaoa, Cupreoboletus, Cacaoporus, Acyanoboletus* and *Cyanoboletus* (Online Resource 2). The taxa in this subgroup were selected to provide the ingroup and outgroup in subsequent phylogenetic reconstructions. Although apparently monophyletic, the species classified in *Cyanoboletus* form a clade with low support (57% posterior probability). In another part of the phylogeny, the placement of *Cyanoboletus flavosanguineus* is compatible with its reclassification as *Neoboletus flavosanguineus* (Biketova et al. 2021), but the phylogenetic tree supporting that proposal cites a doctoral thesis that remains to be made public, so that, to the best of our knowledge, this phylogenetic reconstruction is the first published illustration of its placement.

With this preliminary result at hand, the present work proceeded to sampling exhaustively the related sequences that were found to belong in the ingroup (Materials and Methods). Using examples of *Pulchroboletus, Tropicoboletus* and *Lanmaoa* as outgroup (Gelardi et al. 2023), the ITS region was analyzed in bali-phy (as outlined in Materials and Methods). The resulting phylogenetic tree offers a novel taxonomic topology (Fig. 1), with three well-supported branches for the ingroup sequences, namely one including sequences identified as *Acyanoboletus, Cacaoporus, Cyanoboletus bessettei, Cy. instabilis* and *Cy. fagaceophilus*, a second one for *Cupreoboletus poikilochromus*, and a third one comprising other species of *Cyanoboletus*. These three branches form a clade well-separated from the nearest genus (*Tropicoboletus*) and suggest three well-resolved genera, the first one being named *Cacaoporus* henceforth.

**Fig. 1.**
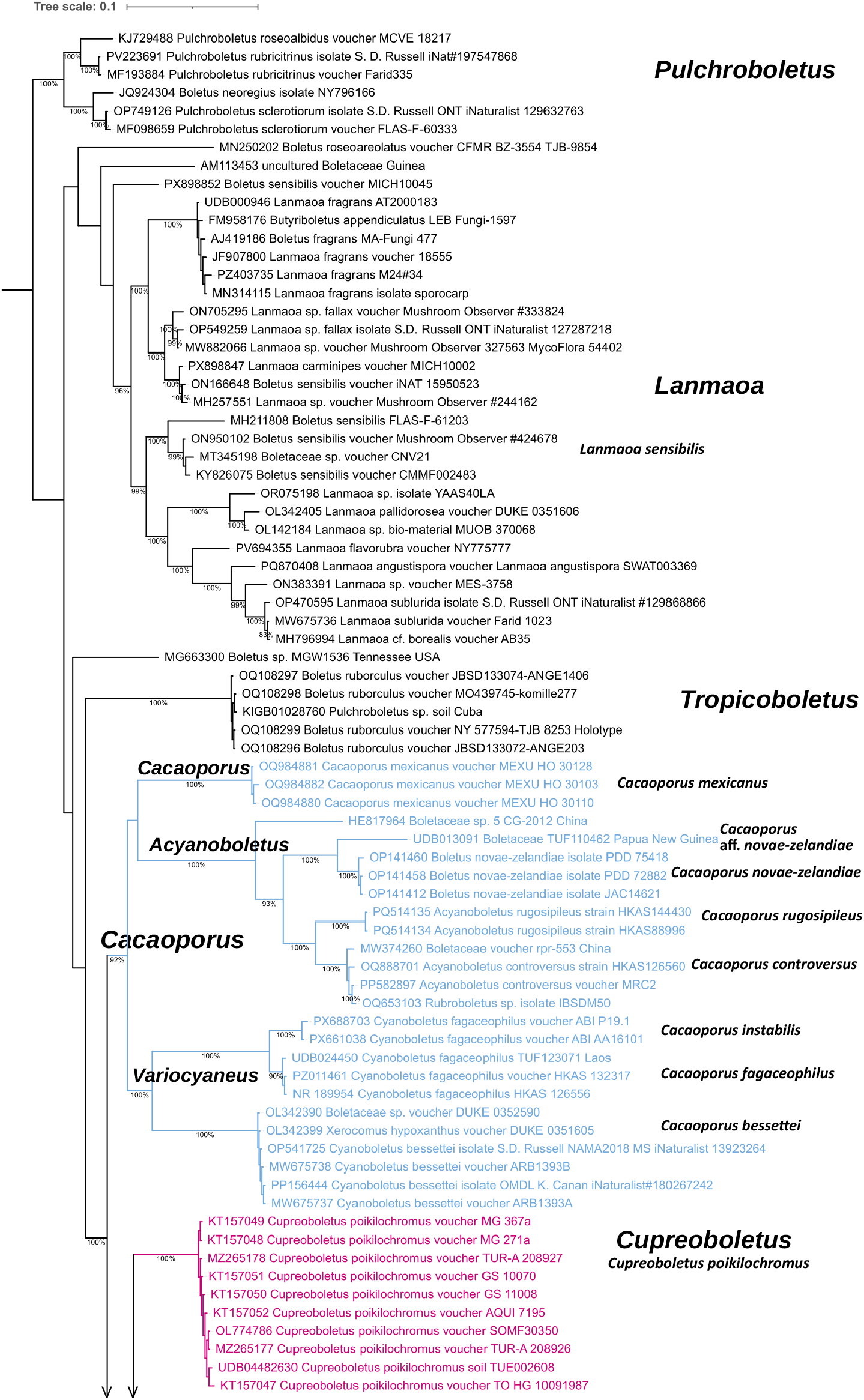
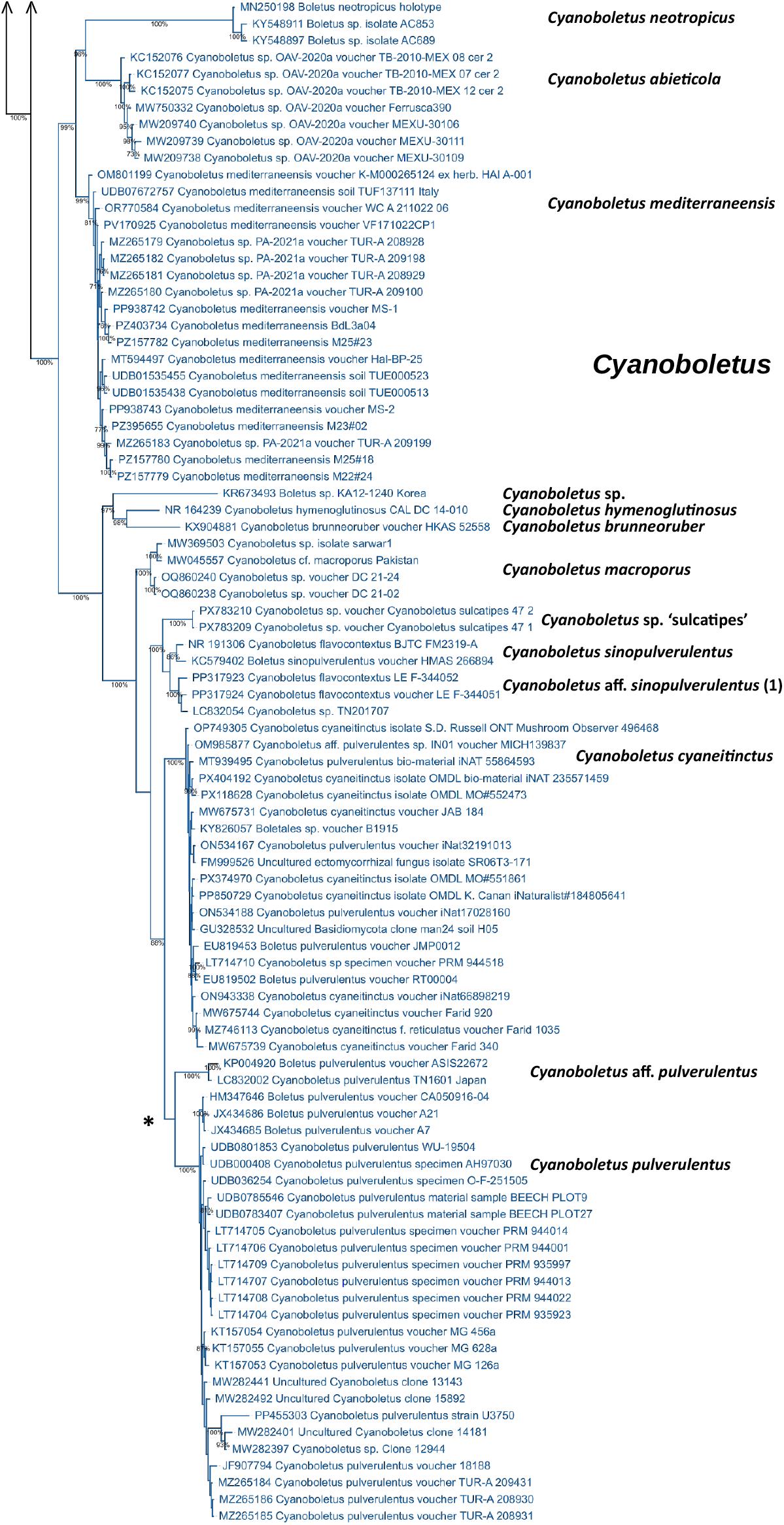
Bali-phy phylogenetic reconstruction for a thorough sampling of clades belonging to *Cyanoboletus* (dark blue), *Cacaoporus* (light blue) and *Cupreoboletus* (purple), based on the nuclear ITS. The MCMC run obtained 17930 samples, the burnin was 1992, and the likelihood ESS was 311. Three redundant leaves were pruned, and node posterior probabilities below 70% are not shown for clarity. Names for clades at genus, subgenus and species-level are indicated in decreasing typeface. Asterisk indicates the ‘*pulverulentus sensu lato*’ clade.

Furthermore, *Boletus novae-zelandiae* McNabb appears in the *Cacaoporus* branch close to the *Acyanoboletus* sequences, and *Boletus neotropicus* B. Ortiz & T.J. Baroni is placed within *Cyanoboletus*. On the other hand, *B. roseoareolatus* B. Ortiz & T.J. Baroni and *B. rugulosiceps* B. Ortiz, T.J. Baroni & Lodge, proposed to belong to *Boletus* sect. *Subpruinosi* along with *B. neotropicus* (Ortiz-Santana et al. 2007), appear not to belong in *Cyanoboletus* and their taxonomic position remains uncertain (Fig. 1, Online Resource 3 Fig. S2a). Most sequences identified as *Boletus sensibilis* Peck suggest that it belongs to genus *Lanmaoa*, although their positions in the *Lanmaoa* clades are diverse.

To explore further taxa without ITS sequences, an ITS+nucLSU concatenate was built, from the nuclear ribosomal RNA large subunit alignment and that of the ITS region with matching specimens. The study on the nucLSU (Online Resource 3) had produced a result completely different from Fig. 1, and this concatenate reflects some of this discrepancy (Online Resource 4) but succeeds in placing at species level most taxa currently represented in the public databases.

To complete the classification of remaining taxa, a concatenate of the nucLSU, RPB2 and TEF1 genes was built, and the resulting phylogeny (Online Resource 5) provided additional placements around *Cacaoporus mexicanus* and *Ca. pallidicarneus*, namely confirming the position of the type species *Ca. tenebrosus*, and a *Cacaoporus* specimen that remains unidentified (SV0402); two further specimens in the *Cyanoboletus* clade (OR0491 and OR0322) are possibly conspecific with *Cyanoboletus brunneoruber* and *Cy. hymenoglutinosus*, respectively (Vadthanarat et al. 2019).

Thus, the following ‘3C’ concept emerges at genus level:

- *Cyanoboletus sensu stricto* encompasses two major branches: one joining *Boletus neotropicus* with *Cyanoboletus abieticola* and *Cy. mediterraneensis*, the other for *Cy. pulverulentus* and neighbouring taxa (*Cy. hymenoglutinosus, Cy. brunneoruber, Cy. macroporus, Cy. viscidiceps, Cy. paurianus, Cy. sinopulverulentus* and *Cy. cyaneitinctus* including its f. *reticulatus*), as well as a minimum of four more taxa, possibly more depending on the putative conspecificity of ‘sulcatipes’ specimens with *Cy. paurianus* (Online Resource 4) and the possible splitting of the East Asian specimens identified as *Cy. pulverulentus* (see below), giving a total of at least 15 species.
- *Cacaoporus* encompasses three major branches: one formed by *Cacaoporus tenebrosus, Ca. pallidicarneus* and *Ca. mexicanus*, and an undescribed collection (SV0402); another by *Acyanoboletus controversus, A. dissimilis* and *A. rugosipileus*, with the addition of *Boletus novae-zelandiae* and three unidentified collections (2 or 3 species); finally, a branch for *Cyanoboletus bessettei, Cy. instabilis, Cy. fagaceophilus* and one more species (collection HKAS 76850), giving a total of at least 14 species. In this new concept of *Cacaoporus*, the three major branches are classified at subgenus level, respectively *Cacaoporus, Acyanoboletus* and *Variocyaneus*.
- *Cupreoboletus*, currently a monospecific genus (*Cu. poikilochromus*).

To investigate the interspecies gap threshold within these three genera, pairwise comparisons of ITS sequence p-distances were calculated, and it was found, by applying increasing values to collapse branches (Online Resource 6), that this gap may start at 0.6% of sequence difference (p-distance). From this result and the phylogenies in Fig. 1 and Online Resources 4 and 5, three cases were singled out for a closer analysis of pairwise p-distances, to clarify the application of correct epithets:

1. The *instabilis*/*fagaceophilus* epithets (*Cacaoporus*). Two clades involve both epithets (Fig. 1, Online Resource 4), the first including specimen HKAS 59554 described as *Cyanoboletus instabilis* (Wu et al. 2016) and the other containing the *fagaceophilus* holotype (specimen HKAS 126556). The net mean p-distance between groups is 3.29%, suggestive of two species (Nilsson et al. 2008; Badotti et al. 2017; Wilson et al. 2023).
2. The *sinopulverulentus*/*flavocontextus* epithets (*Cyanoboletus*). At least three branches contain sequences with these epithets, one featuring the holotypes for both (HMAS 266894 and BJTC FM-2319-A, respectively), a second one that we name ‘aff. *sinopulverulentus* (1)’ and a third one lacking ITS sequences that we name ‘aff. *sinopulverulentus* (2)’. The ITS sequences from the first two branches, when analysed as a single group (*‘sinopulverulentus* s.l., Table 2), show a mean within-group p-distance of 0.99%, suggesting heterogeneity at species level, compared to 0.33% in the group formed by the holotypes and 0.81% for ‘aff. *sinopulverulentus* (1)’. The net between group average p-distance is just at 0.62%. This suggests inferred conspecificity of the two holotypes, such that *Cyanoboletus sinopulverulentus* var. *flavocontextus* is proposed for the more recent holotype in the Taxonomy section. Further analysis of the morphological diagnoses is made in the Discussion.
3. The ‘*pulverulentus* sensu lato’ clade (*Cyanoboletus*). This is the sister group (marked with an asterisk in Fig. 1 and Online Resource 4) to *Cy. cyaneitinctus*; its mean within-group p-distance of 0.92 ± 0.15% prompted the question whether the splits within this clade might represent separate species. Considering four groups (Table 3), the net mean between group p-distances hint at this possibility, especially for accessions KP004920 and LC832002 (‘*pulverulentus* 3’ in Table 3) forming the branch labelled ‘aff. *pulverulentus*’ in Fig. 1.

**Table 2.**
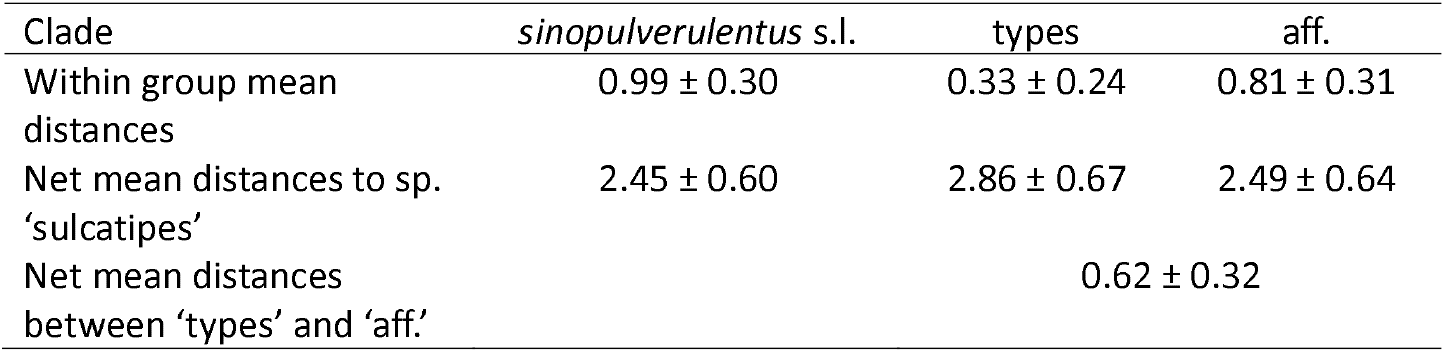
Statistics of pairwise p-distances (ITS, means ± SE, %) for the *Cyanoboletus sinopulverulentus/flavocontextus* pair.

**Table 3.** Statistics of pairwise p-distances (ITS; mean ± SE, %) within and net between groups for the clades close to *Cyanoboletus pulverulentus: pulverulentus* 1 = Cy. *pulverulentus* sensu stricto (24 sequences); *pulverulentus* 2 = accessions MW282397and MW282401; *pulverulentus* 3 = accessions KP004920 and LC832002.

| Clade (within distances) | Net mean distances between groups to clade |  |  | Acc. PP455303 | *cyaneitinctus* |
| --- | --- | --- | --- | --- | --- |
|  | *pulverulentus* 1 | *pulverulentus* 2 | *pulverulentus* 3 |  |  |
| *pulverulentus* 1 (0.07 ± 0.04) | — | 1.16 ± 0.46 | 3.44 ± 0.71 | 3.03 ± 0.11 | 2.61 ± 0.65 |
| *pulverulentus* 2 (0.66 ± 0.32) | | — | 4.30 ± 0.80 | 2.30 ± 0.15 | 3.81 ± 0.79 |
| *pulverulentus* 3 (0.72 ± 0.35) | | | — | 5.89 ± 0.42 | 3.31 ± 0.71 |
| Acc. PP455303 (0) | | | | — | 5.53 ± 0.12 |
| *cyaneitinctus* (0.16 ± 0.05) | | | | | — |

A correspondence of the unidentified clades in Biketova et al. (2026) with our list of clades at species level is presented in Online Resource 7. The current sequence compilation (Online Resource 1) has also allowed a mapping of the occurrences of the two European *Cyanoboletus* species (Online Resource 8).

Finally, to further explore the phylogenetic position of *Neoboletus flavosanguineus* revealed in Online Resource 2, further exploration of close sequences was made. Both the ITS dataset and the LSU+RPB2+TEF1 concatenate (Online Resource 9) suggest that the closest species are represented in a clade named *N. subvelutipes* (Peck) Yang Wang, B. Zhang & Yu Li and a clade encompassing North American collections, namely the holotypes of *Boletus roseobadius* A.H. Sm. & Thiers and *B. rufocinnamomeus* A.H. Sm. & Thiers, as well as collections named *B. subluridellus* A.H. Sm. & Thiers (Tremble et al. 2024) and various other names. The concatenate set further reveals a very close relationship to sequestrate species from Asia, possibly vicariant over the Himalaya: *Gastroboletus thibetanus* Shu R. Wang & Yu Li from China (HKAS 59443) and *Neoboletus angiocarpus* Su. Datta, K. Das & Vizzini from India (KD 24HP-134/142). Of note, *B. rufocinnamomeus* and *B. subluridellus* are reported to develop (microscopic) dextrinoid reactions to Melzer’s reagent (https://www.mushroomexpert.com/boletes_red_pored.html), a feature akin to the amyloid reaction of the context in *N. flavosanguineus*.

## Taxonomy

### Nomenclatural novelties

#### Supraspecific level

*Cacaoporus* Raspé & Vadthanarat subg. *Cacaoporus, subg. nov*. IF XXXXXX. Type species *Cacaoporus tenebrosus* in Vadthanarat, Lumyong & Raspé, *MycoKeys* 54: 20 (2019). Holotype: SV0223 (CMUB)

*Cacaoporus* subg. *Acyanoboletus* Paulo Oliveira & Rodrigo Mariquito, *subg. nov*., *stat. nov*. IF XXXXXX. Type species *Cacaoporus controversus* in Wu, Li, Horak, Wu, Li & Yang, *Mycosphere* 14(1): 753 (2023). Holotype: HKAS 126560.

*Cacaoporus* subg. *Variocyaneus* Paulo Oliveira & Rodrigo Mariquito, *subg. nov*. IF XXXXXX. Type species *Cacaoporus bessettei* in Farid, Bessette, Bessette, Bolin, Kudzma, Franck & Garey, *Mycosphere* 12(1): 1047 (2021). Holotype: ARB1393/USF 301500. Etimology: Latin *vario* = various + *cyaneus* = blue, to denote the diversity of bluish tones that are developed by the different known species on air exposure of the context.

#### Specific level

*Cyanoboletus neotropicus* (B. Ortiz & T.J. Baroni) Paulo Oliveira & Rodrigo Mariquito, *comb. nov*. IF 511050. Basionym: *Boletus neotropicus* B. Ortiz & T.J. Baroni, *Fungal Diversity* 27(2): 312 (2007)

*Cacaoporus bessettei* (A.R. Bessette, Kudzma & Farid) Paulo Oliveira & Rodrigo Mariquito, *comb. nov*. IF 840857. Basionym: *Cyanoboletus bessettei* A.R. Bessette, Kudzma & Farid, in Farid, Bessette, Bessette, Bolin, Kudzma, Franck & Garey, *Mycosphere* 12(1): 1047 (2021)

*Cacaoporus fagaceophilus* (G. Wu, Hai J. Li & Zhu L. Yang) Paulo Oliveira & Rodrigo Mariquito, *comb. nov*. IF 847057. Basionym: *Cyanoboletus fagaceophilus* G. Wu, Hai J. Li & Zhu L. Yang, in Wu, Li, Horak, Wu, Li & Yang, *Mycosphere* 14(1): 759 (2023)

*Cacaoporus instabilis* (W.F. Chiu) Paulo Oliveira & Rodrigo Mariquito, comb. nov. IF 818442. Basionym: *Boletus instabilis* W.F. Chiu, *Mycologia* 40(2): 215 (1948). Synonymy: *Cyanoboletus instabilis* (W.F. Chiu) G. Wu & Zhu L. Yang 2016

*Cacaoporus novae-zelandiae* (McNabb) Paulo Oliveira & Rodrigo Mariquito, *comb. nov*. IF 327059. Basionym: *Boletus novae-zelandiae* McNabb, *New Zealand J. Bot*.: 172 (1968)

*Cacaoporus controversus* (G. Wu & Zhu L. Yang) Paulo Oliveira & Rodrigo Mariquito, *comb. nov*. IF 847055. Basionym: *Acyanoboletus controversus* G. Wu & Zhu L. Yang, in Wu, Li, Horak, Wu, Li & Yang, *Mycosphere* 14(1): 753 (2023)

*Cacaoporus dissimilis* (E. Horak & G. Wu) Paulo Oliveira & Rodrigo Mariquito, *comb. nov*. IF 848543. Basionym: *Acyanoboletus dissimilis* E. Horak & G. Wu, in Wu, Li, Horak, Wu, Li & Yang, *Mycosphere* 14(1): 755 (2023)

*Cacaoporus rugosipileus* (G. Wu & Xin Zhang) Paulo Oliveira & Rodrigo Mariquito, *comb. nov*. IF 855083. Basionym: *Acyanoboletus rugosipileus* G. Wu & Xin Zhang bis, in Zhang, Li, Wang & Wu, *Phytotaxa* 689(2): 208 (2025)

*Lanmaoa sensibilis* (Peck) Paulo Oliveira & Rodrigo Mariquito, *comb. nov*. IF 203533. Basionym: *Boletus sensibilis* Peck, *Ann. Rep. N*.*Y. St. Mus. Nat. Hist*. 32: 33 (1880) [1879]. Synonymy: *Boletus miniato-olivaceus* var. *sensibilis* (Peck) Peck 1889; *Boletus sensibilis* var. *subviscidus* A.H. Sm. & Thiers. 1971

#### Infraspecific level

*Cyanoboletus sinopulverulentus* (Gelardi & Vizzini) Gelardi, Vizzini & Simonini var. *sinopulverulentus* IF XXXXXX. Holotype: HMAS 266894; isotypes TO HG2821 and MG434a in Vizzini, *Index Fungorum* 176: 1 (2014). Basionym: *Boletus sinopulverulentus* Gelardi & Vizzini, *Sydowia* 65(1): 49 (2013). Synonymy: *Cyanoboletus sinopulverulentus* (Gelardi & Vizzini) Gelardi, Vizzini & Simonini 2014

*Cyanoboletus sinopulverulentus* var. *flavocontextus* (L. Fan, N. Mao & T.Y. Zhao) Paulo Oliveira & Rodrigo Mariquito, *comb. & stat. nov*. IF XXXXXX. Holotype: BJTC FM2319-A (Mao). Basionym: *Cyanoboletus flavocontextus* L. Fan, N. Mao & T.Y. Zhao, in Mao, Zhao, Zhang, Li, Lv & Fan, *Mycosphere* 14(1): 2034 (2023)

*Lanmaoa sensibilis* (Peck) var. *sensibilis* Paulo Oliveira & Rodrigo Mariquito, *comb. nov*. IF 426839. Basionym: *Boletus sensibilis* Peck, *Ann. Rep. N*.*Y. St. Mus. Nat. Hist*. 32: 33 (1880) [1879]. Synonymy: *Boletus miniato-olivaceus* var. *sensibilis* (Peck) Peck 1889.

*Lanmaoa sensibilis* var. *subviscidus* (A.H. Sm. & Thiers) Paulo Oliveira & Rodrigo Mariquito, *comb. nov*. IF 347847. Basionym: *Boletus sensibilis* var. *subviscidus* A.H. Sm. & Thiers, *Boletes of Michigan* (Ann Arbor): 286 (1971).

### Taxonomic proposals

#### Description of Cyanoboletus Gelardi, Vizzini & Simonini (Vizzini 2014) emend. Paulo Oliveira & Rodrigo Mariquito

Basidiomes pileate-stipitate with tubular-poroid hymenophore, epigeal, small to medium-small, evelate; pileus tomentose to glabrous, dry to slightly tacky; hymenophore adnate to depressed around the stipe, sometimes decurrent, tubes yellowish, pore surface yellow, orange to reddish, brownish; stipe never ventricose, rarely clavate, surface smooth to pruinose, transversely streaked-scissurate, rarely with a fine reticulum restricted to the stipe apex or with a pseudoreticulate pattern; context yellowish, often reddish-tinged at the base; tissues instantly discolouring dark indigo blue to blue-blackish when handled or injured; taste mild; spore print olive-brown; basidiospores smooth, ellipsoidal to ellipsoidal-fusoid, ellipsoidal-subamygdaliform or narrowly amygdaliform; cystidia cylindrical-fusoid to ventricose-fusoid or lageniform; pileipellis an ixotrichodermium, trichodermium, ixosubcutis or subcutis; hymenophoral trama bilateral-divergent of the ‘Boletus-type’; lateral stipe stratum of the ‘boletoid type’; inamyloid trama in general, dextrinoid in parts of the pileipellis and cystidia in some species; clamp connections absent; ontogenetic development gymnocarpic.

Comment: the original description (Vizzini 2014) was modified by Wu et al. (2016) but not published as an amendment probably because referring to species from China, and it included new characters, foremost being: hymenophore also subdecurrent (as signaled already by Muñoz 2005 for the type species) with a surface sometimes brownish red; pileipellis an ixosubcutis (in line with the ‘tacky’ character above) to subcutis. A formal emendation has been published more recently (Biketova et al. 2026), incorporating these additions but, unfortunately, forcing characters of *Cacaoporus* subg. *Variocyaneus* and *Cupreoboletus* — understandable, given the perceived inclusion of these groups in *Cyanoboletus*, yet with too broad variation — which, according to the present study, should be excluded. Moreover, the statement that the hymenophore surface is “rarely orange, yellowish brown, brownish red to reddish brown” is misleading, since this variation is predominant, not rare, in some *Cyanoboletus* species. On the other hand, the statements in the present emendation on the reticulation of the stipe and basidiospore shape are taken from the study of Biketova et al. (2026).

### Excluded species

*Cyanoboletus pulverulentus* is restored as *Cupreoboletus poikilochromus*, as proposed in the present study.

*Cyanoboletus bessettei, Cy. fagaceophilus* and *Cy. instabilis* are classified in *Cacaoporus* subg. *Variocyaneus*, as proposed in the present study.

*Cyanoboletus flavosanguineus* is a member of genus *Neoboletus* known from Italy (Lavorato and Simonini 1997, Muñoz 2005), as confirmed in the present study (Online Resources 2 and 9), unusual for the genus for having a reticulate stipe and amyloid context. The amyloid reaction may be a character shared with the phylogenetically close North American species *Boletus rufocinnamomeus* and *B. subluridellus*. It is included in the comparison table for European species, presented below (Table 4).

**Table 4.** Diagnosis of European taxa.

| Species | *Cupreoboletus poikilochromus*<br>(Muñoz 2005; Gelardi et al. 2015) | *Cyanoboletus mediterraneensis*<br>(Biketova et al. 2022; unpublished observations) | *Cyanoboletus pulverulentus*<br>(Muñoz 2005) | *Neoboletus flavosanguineus*<br>(Muñoz 2005) |
| --- | --- | --- | --- | --- |
| Pileus surface | Viscid in moist weather | Viscidulous | Dry | Dry |
| Hymenophore surface | Yellow, later copper brown | Yellow, later olivaceous and rusty orange | Yellow, later olive yellow | Yellow, later reddish |
| Stipe shape | Initially ventricose, then (sub)cylindrical | Cylindrical to fusiform | Cylindrical, broadening at the apex | Initially slightly ventricose, then slightly clavate towards the base |
| Stipe reticulum | Well-developed, at least top third | Close to the hymenophore, sometimes absent | Close to the hymenophore, frequently absent | Extending only over the top, sometimes down to the whole stipe |
| Context | Probably inamyloid | Inamyloid | Inamyloid | Strongly amyloid |
| Odour | Strong and persistent, of propolis, fermented fruit, etc. | Inconspicuous (slightly acidulous/fruity) | Inconspicuous (slightly acidulous/fruity) | Inconspicuous (slightly acidulous/fruity) |
| Basidiospore dimensions | Length <12.5 µm, width < 5.5 µm, $Q_m < 2.45$ | Width $\geq 4.8$ µm, $Q_m < 2.45$ | Width < 5.5 µm, $Q_m > 2.45$ | Length $\geq 12.5$ µm, width > 5.5 µm, $Q_m < 2.45$ |
| Suprahilar depression<sup>a</sup> | Poorly defined | Occasionally obvious (generally < 2.5% of convex hull area) | Pronounced in a large proportion of spores (generally > 3% of convex hull area) | Poorly defined |
| Gelatinised pileipellis | Yes | Yes | No | No |
| Encrustations in pileipellis | Rare, subtle | Intracellular | Intracellular | None |

<sup>a</sup> Based on Biketova et al. (2026); for *N. flavosanguineus*, by analogy, based on Lavorato and Simonini (1997).

*Cyanoboletus rainisiae* is a North American member of genus *Xerocomellus* (Frank et al. 2020), with dark bluish staining of the pileus, stipe and hymenophore surface where touched, as well as a dark blue staining in all parts of the exposed context, characters that are quite unusual for the genus and have led to confusions with *Boletus pulverulentus* (currently known as *Cyanoboletus cyaneitinctus*) and its initial inclusion in *Cyanoboletus* (Vizzini 2014). However, the areolate pileus, especially at the margin, common with maturation, is shared with many species of *Xerocomellus* and does not fit the description of *Cyanoboletus*. Given its taxonomic placement, one should expect a hymenophoral trama intermediate between the phylloporoid and boletoid types, with lateral strata weakly but distinctly gelatinous, especially in younger basidiomes (Šutara 2008), while in *Cyanoboletus* it is of the boletoid type.

*Cyanoboletus gabretae* is a seldom described species, originally compared to *B. junquilleus* (Quél.) Costantin & L.M. Dufour but with a well-developed reticulum on the stipe (Pilát 1968; Kallio 1984), also with an overall appearance comparable to xanthoid variants of *Suillellus luridus* (Schaeff.) Murrill (Muñoz 2005), in spite of the different type of reticulum (Kallio 1984) or, better still, as a smaller, conifer-associated lookalike of xanthoid forms of genus *Imperator* (Klofac and Krisai-Greilhuber 2018). It does not fit the description of *Cyanoboletus* because of the persistently yellow pileus (in maturity less bright, but without shades of brown) and the reticulate stipe, ventricose in young specimens. Further discussion of its taxonomic placement is made by Biketova et al. (2026).

#### Description of Cacaoporus Raspé & Vadthanarat ex Paulo Oliveira & Rodrigo Mariquito

Basidiomata stipitate-pileate with poroid hymenophore, small to medium-sized. Pileus convex with inrolled margin when young, becoming plano-convex to slightly depressed with age, with deflexed to inflexed margin; surface even to rugulose, nearly glabrous to subtomentose, sometimes slightly cracked at the centre. Hymenophore tubulate, adnate to subdecurrent, tubes pale yellow or brownish; pores concolorous, regularly arranged, mostly roundish at first and becoming slightly angular with age, sometimes irregular, elongated around the stipe. Stipe central, cylindrical, may be enlarged or tapered at the base; surface even, minutely tomentose. Spore print dark brown or olive-brown. Context pale yellowish to greyish off-white, sometimes with scattered small dark brown to brownish-black encrustations, sometimes changing immediately to reddish or blue-green on exposure to air, sometimes with delayed colouring. Basidiospores subfusoid, amygdaliform, ellipsoid or ovoid, thin-walled, smooth. Pileipellis a more or less tangled trichoderm. Clamp connections absent.

#### Diagnosis of Cupreoboletus (Muñoz (2005), updated with Gelardi et al. (2015) and Biketova et al. (2026))

Differs from *Cyanoboletus* and *Cacaoporus* by the stipe with well-developed, wide-mesh, longitudinally stretched reticulum extending over the upper half, stipe frequently clavate, even ventricose in immature basidiomes, the exposed context turning cinnamon orange/orange red after some hours, and strong sweetish odour reminiscent of propolis, liqueur, fermented fruits, dried cinnamon or cottonwood.

#### Diagnosis of Cacaoporus

Differs from *Cyanoboletus* by the subtomentose to minutely velutinous pileus surface, that may become glabrous, in some species more or less rugulose and/or cracking in the disc, reactions to bruising or exposure to air diverse or absent, never the typical deep ultramarine blue oxidation of *Cyanoboletus*, odour frequently distinct and unpleasant, pileipellis a trichoderm or subcutis, not gelatinised, hymenophoral trama of the boletoid or sometimes intermediate type.

We note that the undescribed specimen HKAS 76850 (*Cacaoporus* aff. *bessettei*, Fig. 3) is mentioned as having a ‘viscid basidioma’ (Wu et al. 2016), therein foreshadowing a possible adjustment of this diagnosis once a proper description of the relevant species is available.

#### Diagnosis of subgenera of Cacaoporus

*Cacaoporus* subg. *Cacaoporus*: basidiome dull, brown to greyish-brown to dark brown or blackish-brown, context pale grey to pale yellow, darkened with marmorated or virgated brown, developing a reddish to greyish-brown reaction on exposure, especially in the pileus; tubes brown to greyish-brown to dark brown, not separable from the pileus context. Basal mycelium (stipe) becoming reddish-white to pale red or to violet–brown when touched. Hymenial cystidia fusiform, cylindrical, clavate, ventricose-rostrate, with obtuse apex.

*Cacaoporus* subg. *Acyanoboletus*: context pale yellow, unchanging on exposure, sometimes developing a delayed colour reaction; hymenophoral surface and tubes pale yellow, concolorous, unchanging when bruised. Stipe yellowish, basal mycelium unchanging when touched. Hymenial cystidia lanceolate to narrowly fusoid, with long or short beaks, or cylindrical with obtuse apex.

*Cacaoporus* subg. *Variocyaneus*: context pale yellow, changing immediately to pale blue/blue-green on exposure, sometimes further developing a different, delayed colour reaction. Stipe with pale pinkish to reddish-brown tone, basal mycelium unchanging when touched. Hymenial cystidia narrowly fusoid to ventricose-fusoid with long or short beaks.

#### Comparison tables

Biketova et al. (2026) note the critical discrimination between *Cyanoboletus mediterraneensis* and *Cy. pulverulentus*, detailing the subtle differences that can be useful, especially in combination, to separate them. Apart from the basidiospore dimensions and suprahilar depression, further distinctions between these two species are set out in Table 4, based on observations of *Cy. mediterraneensis* specimens from South Portugal (to be published), and may or may not be confirmed with their study in other materials. For example, the evolution of the hymenophore surface to rusty orange is a very common character in the Portuguese collections studied by us, as is the pileipellis, gelatinised in *Cy. mediterraneensis* and non-gelatinised in *Cy. pulverulentus* (Muñoz 2005). The fusiform stipe can be seen in several photos of *Cy. mediterraneensis* in Biketova et al. (2026) and can be an accessory character of interest.

**Table 5.**
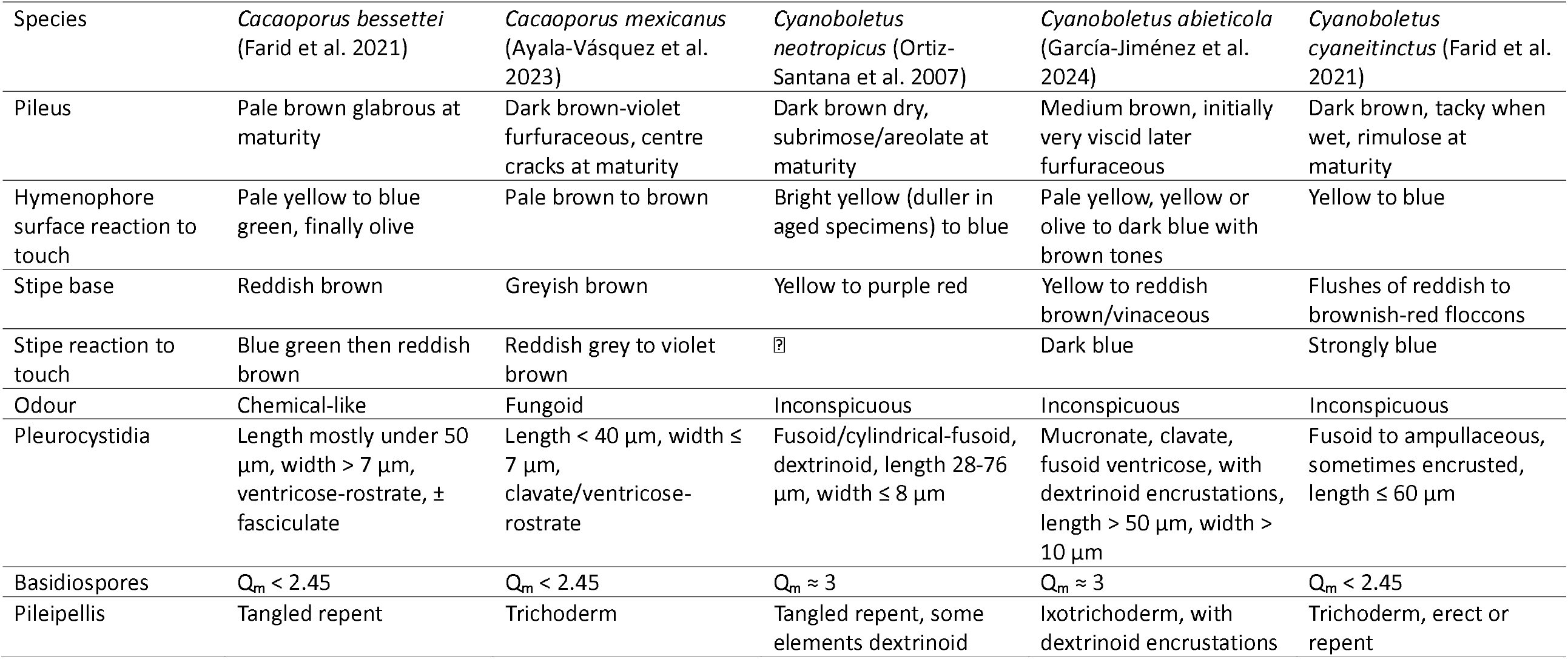
Diagnosis of American taxa.

#### Key to Asian-Oceanic species of Cacaoporus

1. Context pale grey-yellow or dark brown, darkened with marmorated brown, developing orange or reddish on exposure, basal mycelium turns reddish-white to pale red when bruised, pileus surface sometimes cracked in the disc, taste slightly bitter at first, average spore quotient Q_m_ < 1.7, cystidia and basidia covered with dark particles or crystals (Subg. *Cacaoporus*) 2 — Context pale yellow, not marmorated, unchanging or with immediate reaction to exposure, later developing a delayed reaction or not, average spore quotient Q_m_ ≥2. 3
2. Pileus context yellowish to greyish off-white, then slightly pale orange to greyish-orange when exposed, odour rubbery, spores amygdaliform *Cacaoporus pallidicarneus* — Pileus context greyish-brown to dark brown, slightly reddening in paler spots when cut, odour mild fungoid, spores ovoid *Cacaoporus tenebrosus*
3. Hymenophore staining blue or blue-green when bruised, stipe surface usually subconcolorous with the pileus or with a pinkish to reddish-brown tone(Subg. *Variocyaneus*) 4 — Hymenophore unchanged when bruised, stipe surface frequently of distinct yellow colour …………………………………………………………………………………………………… (Subg. *Acyanoboletus*) 5
4. Stipe context changing to blue on exposure, pileipellis a subcutis *Cacaoporus instabilis* — Stipe context almost unchanging on exposure, pileipellis a trichoderm ………………………………………………………………………………………………… *Cacaoporus fagaceophilus*
5. Pileus surface rugose at maturity, especially towards the margin, slightly viscid when wet, pileipellis an intricate trichoderm to interwoven subcutis *Cacaoporus rugosipileus* — Pileus surface smooth, felted to subtomentose at least initially, pileipellis a trichoderm 6
6. Stipe remarkably shorter than the pileus diameter, basidiome colours yellowish, basidiospore length > 14 μm, cystidia (sub)cylindrical *Cacaoporus novae-zelandiae* — Stipe approximately the same length as the pileus diameter, basidiome colours diverse, basidiospore length < 14 μm, pleurocystidia lanceolate to fusoid 7
7. Pileus greyish-brownish orange, without red tinges, odour of coal gas, basidiospore length < 11 μm *Cacaoporus controversus* — Pileus pale chestnut brown, more or less reddish, odour of pharmacy, basidiospore length > 11 μm *Cacaoporus dissimilis*

#### Key to Asian species of Cyanoboletus

1. Hymenophore surface bright yellow, pileus without red/orange tinge, typically viscid when wet, 2-spored basidia present 2 — Hymenophore surface predominately orange to brown, pileus brown, brownish orange or brownish red, stipe dark reddish brown 5
2. Pileus light brown, stipe deep red downwards, average spore quotient Q_m_ > 2.45 ……………………………………………………………………………………………………*Cyanoboletus viscidiceps* — Pileus dark brown 3
3. Stipe evenly dark brown, with brownish-pink shades at the base, finely transversely streaked-scissurate (zebroid pattern) in the upper half, context whitish especially in the pileus, basidia mainly 2-spored, average spore quotient Q_m_ < 2.45 ……………………………………………………….. *Cyanoboletus sinopulverulentus* var. *sinopulverulentus* — Stipe yellow, brown towards the base, surface glabrous to leathery, basidia mainly 4-spored …………………………………………………………………………………………………………………………………….. 4
4. Pleurocystidia up to 35 μm long, spore quotient 1.65–2.86 (Q_m_ < 2.45) …………………………………………………………………………………………………… *Cyanoboletus paurianus* — Pleurocystidia at least 35 μm long, spore quotient 2.18–2.89 (Q_m_ > 2.45) …………………………………………………………….*Cyanoboletus sinopulverulentus* var. *flavocontextus*
5. Glutinous covering of all surfaces including the hymenium, tube pores initially stuffed, 2-spored basidia frequent, pleurocystidia mostly < 50 μm long *Cyanoboletus hymenoglutinosus* — Non-glutinous, tube pores not stuffed, 2-spore basidia absent or rare, pleurocystidia length mostly > 50 μm 6
6. Hymenophore with wide pores (<1/mm), habitat temperate to subalpine ………………………………………………………………………………………………… *Cyanoboletus macroporus* — Hymenophore with narrow pores (>1/mm), habitat subtropical *Cyanoboletus brunneoruber*

## Discussion

When this work was started in 2025, chiefly to study the complex of species under the name *Cyanoboletus pulverulentus* and to ascertain the taxonomic position of the materials collected in the Alentejo region in Portugal, the authors did not have any hint of how muddled the taxonomic situation was. The genera *Cyanoboletus* (Vizzini 2014) and *Cupreoboletus* (Gelardi et al. 2015) had been created from previously known taxa in 2014 and 2015 respectively, then several atypical (staining-wise) “basal” species were integrated in the former (Wu et al. 2016; Farid et al. 2021; Wu et al. 2023) and *Cupreoboletus* ended up in *Cyanoboletus* as well (Carbone et al. 2023). Such dumping onto *Cyanoboletus* introduced increasingly dramatic contradictions with the genus diagnosis, while the alternative of recognizing diverging genera only came by to address the particular morphology of *Cacaoporus* (Vadthanarat et al. 2019) and the absence of staining reactions of *Acyanoboletus* (Wu et al. 2023).

The ITS alignment and resulting phylogeny that we obtained with bali-phy (Redelings 2021) challenges the current taxonomy, and merits consideration in view of the unprecedented clarity of taxon relationships at both molecular and morphological levels. The main cause for the discrepancies between gene datasets lies in the highly contrasting divergence rates of the nucLSU and protein coding genes between the three subgenera of the now revised *Cacaoporus*, to the effect of dispersing them in the multigene tree, even “enveloping” *Cupreoboletus* (Online Resources 2, 3 and 5). A comparison of the single-gene phylogenies (Online Resource 3) reveals glaring contradictions, making the concatenations of these genes— the only molecular approach that has been used to address evolutionary relationships within this group—highly doubtful. In other words, we suspect that those markers, so important to outline higher-order phylogenetic reconstructions and place the root in our trees (Gelardi et al. 2023), are unsuitable at this level. By contrast, the evolutionarily neutral variation in the ITS is expected to render the most faithful representation of the macroevolutionary process (Kimura 1983, Nei and Kumar 2000), provided that a reliable alignment is achieved, and a sign that this was successful is the greater homogeneity of branch lengths among clades (Fig. 1).

The usefulness of the ITS marker for phylogenetic studies of divergent taxa groups has been hampered by difficulties in objectively aligning sequences that contain numerous indels. However, the bali-phy toolset (Redelings 2021) proposes a solution to manage this, and the experience in the present study was remarkably powerful. Especially with high computing power to reduce response times, it should be a worthwhile endeavor to find out just how far apart the divergence between sequences can be tackled with it.

The epithets *poikilochromus, bessettei, fagaceophilus* and *instabilis*, if kept in *Cyanoboletus*, demand a significant change in the emendation of this genus diagnosis (Biketova et al. 2026); conversely, the present work suggests that *Cupreoboletus poikilochromus*, while branching close to *Cyanoboletus*, is not a morphological and organoleptic quirk in the diversification of the latter, but a separate genus; this, together with the unsuspected unification of the other three in *Cacaoporus* restores a coherent, distinct, useful concept of *Cyanoboletus*. We surmise, the mentioned emendation should not be adopted, and a description that maintains the original concept of *Cyanoboletus*, updating a few details, is proposed here instead (Taxonomy section).

Unlike the mounting problems that were happening with the delimitation of *Cyanoboletus*, the current proposal of *Cacaoporus* as the home for *Acyanoboletus* and the species now forming subgenus *Variocyaneus* (Fig. 1) does not create any difficulties; quite the reverse, it is supported by a fairly simple diagnosis of the genus and its subgenera. Incidentally, the discovery that *Boletus novae-zelandiae* belongs in *Cacaoporus* subg. *Acyanoboletus* contributes to a more nuanced appreciation of the color changes of the context and of the cystidia shapes for this subgenus (McNabb 1968).

One important application of the proposed taxonomy is to instruct future phylogenetic studies that will include the ‘3C’ genera, namely, to strategize taxon sampling outlooks that make for a balanced representation of the main clades, *viz*. at least one taxon for each subgenus of *Cacaoporus*, at least one taxon for each of the two longer branches within *Cyanoboletus*, and *Cupreoboletus* (Fig. 1).

Our analysis highlights a critical need for attention to the usage of some epithets:

i. The epithets *instabilis* and *fagaceophilus* (*Cacaoporus*). From the protologues (Wu et al. 2016, Wu et al. 2023) there is some evidence in favour of considering these as separate species, especially for the differences in the pileipellis structure, possibly aided by a difference in the bluing of the context on exposure that would require a more precise characterization. In the phylogeny there are two clades with these two epithets (Fig. 1) separated by a sizeable mean p-distance of 3.29%, strongly supporting the separation of two species, and indicating that at least some of the sequenced specimens have a wrong epithet. In contrast to the *fagaceophilus* holotype (voucher HKAS 126556) placed in one of these clades, the position of the *instabilis* holotype is still unknown; the other clade contains the HKAS 59554 specimen included in a description of *Cyanoboletus instabilis* (Wu et al. 2016), thus encouraging naming this clade with the *instabilis* epithet (Fig. 1, Online Resources 4 and 6). But the uncertainties point out the need for a deeper understanding of the diagnosis between the two species, and for sequences from a *Cy. instabilis* type, while cautioning on the application of these epithets until a more grounded clarification is achieved.
ii. The epithets *sinopulverulentus*/*flavocontextus* (*Cyanoboletus*). The average p-distance of 0.33% within the group consisting of the respective holotypes (both from the Shaanxi province, China) would suggest conspecificity (Table 2). Comparison of the descriptions in the protologues (Gelardi et al. 2013, Mao et al. 2023) does not bring forth substantive differences except for the texture and colours of the stipe, and also the context colour, described as “pale yellow” (pileus) and “bright yellow, turning orange-yellow from bottom up with age” (stipe) for *Cy. flavocontextus* (Mao et al. 2023), in contrast to the “whitish” (pileus) and “very pale yellow” (stipe) for *Cy. sinopulverulentus* (Gelardi et al. 2013). On the other hand, the description of *Cy. sinopulverulentus* by Wu et al. (2016), citing the holotype as well as a sequenced specimen from Yunnan, China (HKAS 59609) that clusters away in the ‘aff. *sinopulverulentus* (2)’ group (Online Resources 4 and 6, close but probably not conspecific to a collection from India, DC 16-51) states “yellowish to yellow” (pileus) and “yellow to orange” (stipe). The present analysis suggests that specimens identified as *Cy. sinopulverulentus* or *Cy. flavocontextus* are part of a species complex that requires further taxonomic study which will also benefit from the inclusion of future materials, so that the diagnosis pertaining to these epithets is settled. It is likely that more species in this complex will emerge, most probably from the two ‘aff. *sinopulverulentus*’ clades defined herein. Based on the putative conspecificiy of the two types above, the holotype of *Cyanoboletus flavocontextus* is reclassified as a variety of *C. sinopulverulentus, Cyanoboletus sinopulverulentus* var. *flavocontextus* (L. Fan, N. Mao & T.Y. Zhao) Paulo Oliveira & Rodrigo Mariquito.

Furthermore, the delimitation of *Cyanoboletus pulverulentus* also needs further work. A group of two Asian sequences branching away (‘aff. *pulverulentus’* in Fig. 1 and Online Resource 4, ‘*pulverulentus* 3’ in Table 3) are as separate from the main ‘*pulverulentus* 1’ group as *Cy. cyaneitinctus* (Table 3). This underscores the need for a thorough study of the corresponding vouchers to achieve a workable diagnosis (other than geography) between them and *Cy. pulverulentus sensu stricto*. The other groups being considered in Table 3 are two ectomycorrhizal sequences from Spain (‘*pulverulentus* 2’) and an accession from Sweden (PP455303), but since they cluster inside the group represented by ‘*pulverulentus* 1’ (Fig. 1), it seems less likely that they are from separate species.

## Conclusion

Results presented in this study argue for a concept of the Suillelloideae genera *Cupreoboletus, Cacaoporus* and *Cyanoboletus* that markedly deviates from the preceding taxonomy and is consistent with well-delimited morphological diagnoses. The proposed regional diagnoses and keys should help a practical and effective determination of new materials. The phylogenetic reconstruction in Fig. 1 provides a framework for a balanced taxon sampling in future studies and, together with Online Resources 4 and 5, points out undescribed taxa as well as taxa still needing a firmer diagnosis. The approach to sequence alignment made by bali-phy encourages tackling the difficulties with the ITS region in broader evolutionary frames.

## Acknowledgements

The authors declare that no funds, grants, or other support were received during the preparation of this manuscript.

## Statements and Declarations

### Author Contributions

Both authors participated in conception and design, sequence compilation, preparation of alignments and phylogenetic analyses. The manuscript draft was prepared by Paulo Oliveira and both authors revised it, read and approved its final form.

### Data Availability

Alignment FastA files, BEAUti XML files, and NEXUS tree files, as well as the bali-phy output, are available in a Figshare repository doi: 10.6084/m9.figshare.32298045.

## Graphical abstract

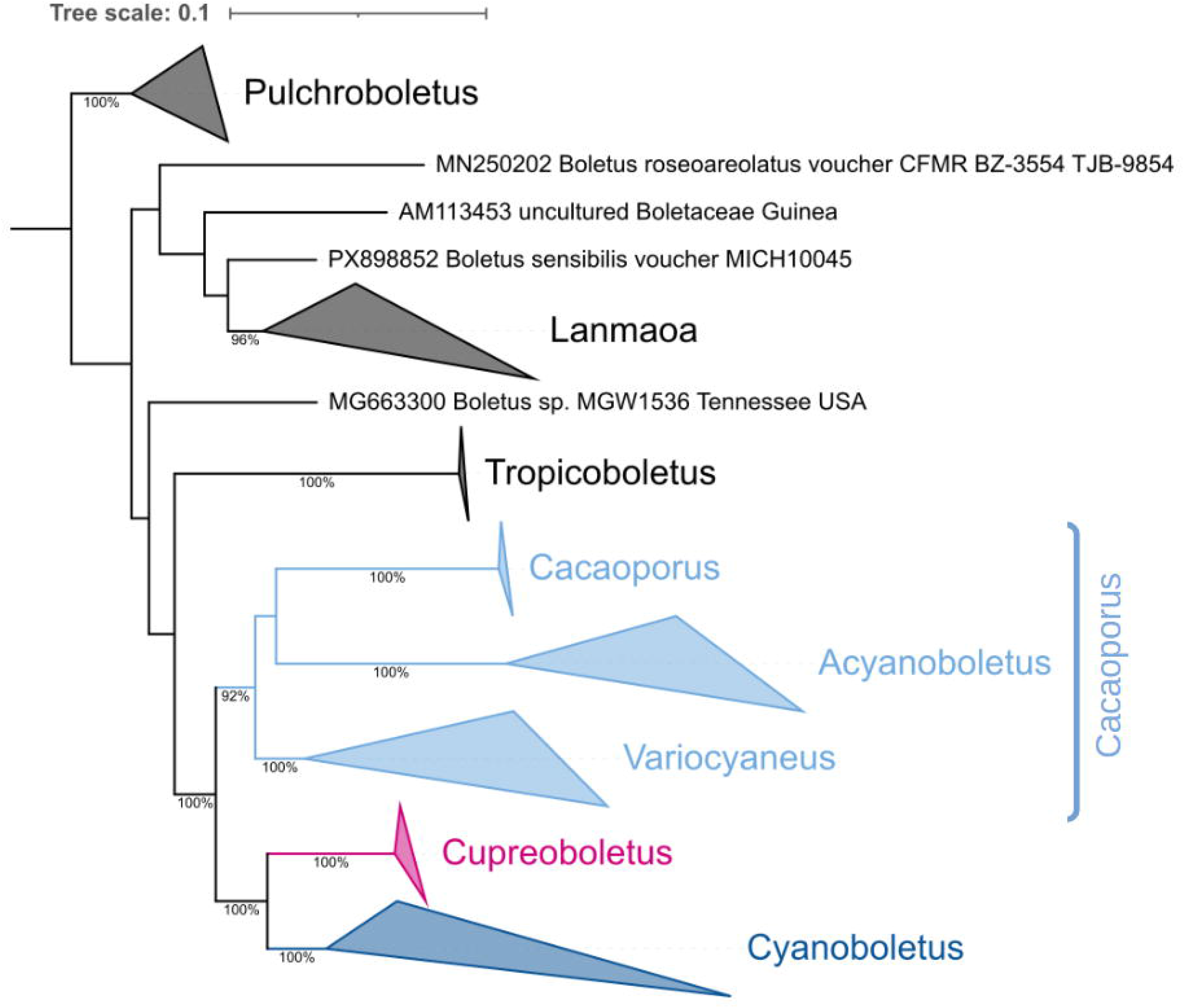

