## Supplementary tables and figures, and legends for "Global delimitation of *Cyanoboletus, Cacaoporus* and *Cupreoboletus* (Suillelloideae : Boletaceae)"

*Mycological Progress* ISSN 1617-416X

Paulo Oliveira[1, 2, 3], Rodrigo Mariquito[1]

[1] Biology Department, University of Évora, Évora, Portugal

[2] Mediterranean Institute for Agriculture, Environment and Development (MED), Évora, Portugal

[3] Research Centre in Biodiversity (CIBIO), Vairão, Portugal

#### Contents

Online Resource 1: Tables S1–S3 (sequence accession codes)

Online Resource 2: Figure S1 (Suillelloidae four-gene phylogeny)

Online Resource 3: Figure S2 (single-gene phylogenies: nucLSU, RPB1, RPB2 and TEF1)

Online Resource 4: Figure S3 (ITS + nucLSU concatenate phylogeny)

Online Resource 5: Figure S4 (nucLSU+RPB2+TEF1 concatenate phylogeny)

Online Resource 6: Figure S5 (pairwise Minimum Evolution ITS phylogeny)

Online Resource 7: Supplementary Table S4 (unidentified clades)

Online Resource 8: Figure S6 (geographic distribution of the two species of European *Cyanoboletus*)

Online Resource 9: Figure S7 (*Neoboletus flavosanguineus* study)

Online Resource 10: Supplementary Figure Legends

### Online Resource 1

Excluded accessions: PV695610, KF668326, KF739691 (redundant); JN563871, JN563872, KF030364, MH645599 (outside the scope).

#### Table S1. List of Genbank accession numbers for the ingroup sequences

| Genus | Species clade | Specimen | Country | ITS | LSU | RPB1 | RPB2 | TEF1 |
|---|---|---|---|---|---|---|---|---|
| Cacaoporus | bessettei | Boletaceae sp. voucher DUKE:0352590 | USA | OL342390 | | | | |
| | | Cyanoboletus bessettei isolate OMDL iNat # 180267242 | USA | PP156444 | | | | |
| | | Cyanoboletus bessettei isolate S.D. Russell NAMA2018 MS iNaturalist 13923264 | USA | OP541725 | | | | |
| | | Cyanoboletus bessettei voucher ARB1393A | USA | MW675737 | MW662571 | | MW737457 | MW737482 |
| | | Cyanoboletus bessettei voucher ARB1393B | USA | MW675738 | | | MW737458 | MW737483 |
| | | Cyanoboletus bessettei voucher MISS0098810 | USA | PV694398 | | | | |
| | | Xerocomus hypoxanthus voucher DUKE:0351605 | USA | OL342399 | | | | |
| | bessettei aff. | Boletus sp. HKAS 76850 | China | | KF112343 | KF112527 | KF112697 | KF112187 |
| | controversus | Acyanoboletus controversus HKAS 126560 | China | NR_189953 | NG_229118 | OQ873469 | OQ873490 | OQ873451 |
| | | Acyanoboletus controversus strain HKAS101248 | China | | OQ888715 | OQ873470 | OQ873491 | OQ873452 |
| | | Boletaceae sp. voucher rpr-553 | China | MW374260 | | | | |
| | | Boletus sp. HKAS 50292 | China | | KF112470 | | KF112816 | |
| | | Acyanoboletus controversus voucher MRC2 | India | PP582897 | | | | |
| | | Rubroboletus sp. isolate IBSDM50 | India | OQ653103 | | | | |
| | dissimilis | Acyanoboletus dissimilis ZT 14030 | Malaysia | | NG_242127 | OQ873471 | OQ873492 | OQ873453 |
| | fagaceophilus | Cyanoboletus fagaceophilus HKAS 126556 | China | NR_189954 | NG_229119 | OQ873473 | OQ873494 | OQ873455 |
| | | Cyanoboletus fagaceophilus strain HKAS80691 | China | | OQ888719 | OQ873474 | OQ873495 | OQ873456 |
| | | Cyanoboletus fagaceophilus voucher HKAS:132317 | China | PZ011461 | | | | |
| | | Cyanoboletus fagaceophilus voucher HKAS123872 | China | | OQ888717 | OQ873472 | OQ873493 | OQ873454 |
| | | Cyanoboletus instabilis voucher N.K.Zeng2862 | China | MG030473 | MG030466 | | | MG030478 |
| | | Cyanoboletus fagaceophilus TUF123071 | Laos | UDB024450 | | | | |
| | instabilis | Boletus sp. HKAS 59554 | China | | KF112412 | KF112528 | KF112698 | KF112186 |
| | | Cyanoboletus fagaceophilus voucher ABI AA16101 | Viet Nam | PX661038 | | | | |
| | | Cyanoboletus fagaceophilus voucher ABI-P19.1 | Viet Nam | PX688703 | | | | |
| | mexicanus | Cacaoporus mexicanus voucher ITCV:1027 | Mexico | | OQ975747 | | OQ938896 | |
| | | Cacaoporus mexicanus voucher ITCV:22674 | Mexico | | | | OQ938892 | |
| | | Cacaoporus mexicanus voucher MEXU:HO 30103 | Mexico | OQ984882 | OQ975750 | | OQ938899 | |
| | | Cacaoporus mexicanus voucher MEXU:HO 30110 | Mexico | OQ984880 | OQ975748 | | OQ938897 | |

| Genus | Species clade | Specimen | Country | ITS | LSU | RPB1 | RPB2 | TEF1 |
|---|---|---|---|---|---|---|---|---|
| | | Cacaoporus mexicanus voucher MEXU:HO 30128 | Mexico | OQ984881 | OQ975749 | | OQ938898 | |
| | novae-zelandiae | Boletus novae-zelandiae isolate JAC14621 | New Zealand | OP141412 | | | | |
| | | Boletus novae-zelandiae isolate PDD 72882 | New Zealand | OP141458 | OP141580 | | | |
| | | Boletus novae-zelandiae isolate PDD 75418 | New Zealand | OP141460 | OP141582 | | | |
| | pallidicarneus | Cacaoporus pallidicarneus HKAS 52601 | China | | KF112469 | KF112552 | KF112732 | |
| | | Cacaoporus pallidicarneus voucher OR0681 | Thailand | | | | MK372283 | |
| | | Cacaoporus pallidicarneus voucher OR0683 | Thailand | | | | MK372284 | |
| | | Cacaoporus pallidicarneus voucher OR1306 | Thailand | | | | MK372285 | MK372272 |
| | | Cacaoporus pallidicarneus voucher SV0221 | Thailand | | | | MK372286 | MK372273 |
| | | Cacaoporus pallidicarneus voucher SV0451 | Thailand | | | | MK372287 | MK372274 |
| | rugosipileus | Acyanoboletus rugosipileus HKAS136929 | China | | PQ517513 | PQ536879 | PQ536883 | PQ536873 |
| | | Acyanoboletus rugosipileus HKAS144429 | China | | PQ517516 | PQ536881 | PQ536886 | PQ536876 |
| | | Acyanoboletus rugosipileus HKAS144430 | China | PQ514135 | PQ517514 | PQ536878 | PQ536885 | PQ536875 |
| | | Acyanoboletus rugosipileus HKAS144431 | China | | PQ517515 | PQ536880 | PQ536884 | PQ536874 |
| | | Acyanoboletus rugosipileus HKAS88996 | China | PQ514134 | PQ517512 | PQ536877 | PQ536882 | PQ536872 |
| | sp. | Boletaceae sp. 5 CG-2012 (subg. *Acyanoboletus*) | China | HE817964 | | | | |
| | | Boletaceae TUF110462 (subg. *Acyanoboletus*) | Papua New Guinea | UDB013091 | | | | |
| | | Cacaoporus sp. voucher SV0402 (subg. *Cacaoporus*) | Thailand | | | | MK372293 | MK372281 |
| | | Boletus sp. 4696 (subg. *Acyanoboletus*) | ? | | KF030331 | | | |
| | tenebrosus | Cacaoporus tenebrosus voucher OR0654 | Thailand | | | | MK372288 | MK372275 |
| | | Cacaoporus tenebrosus voucher OR1435 | Thailand | | | | MK372289 | MK372276 |
| | | Cacaoporus tenebrosus voucher SV0223 | Thailand | | | | MK372290 | MK372277 |
| | | Cacaoporus tenebrosus voucher SV0224 | Thailand | | | | MK372291 | MK372278 |
| | | Cacaoporus tenebrosus voucher SV0422 | Thailand | | | | | MK372279 |
| | | Cacaoporus tenebrosus voucher SV0452 | Thailand | | | | MK372292 | MK372280 |
| Cupreoboletus | poikilochromus | Cupreoboletus poikilochromus voucher SOMF30350 | Bulgaria | OL774786 | | | | |
| | | Cupreoboletus poikilochromus soil TUE002608 | Italy | UDB04482630 | | | | |
| | | Cupreoboletus poikilochromus voucher AQUI 7195 | Italy | KT157052 | KT157061 | | | |
| | | Cupreoboletus poikilochromus voucher GS 10070 | Italy | KT157051 | KT157060 | KT157066 | KT157068 | KT157072 |
| | | Cupreoboletus poikilochromus voucher GS 11008 | Italy | KT157050 | KT157059 | KT157065 | KT157067 | KT157071 |
| | | Cupreoboletus poikilochromus voucher MG 271a | Italy | KT157048 | KT157057 | | | KT157070 |
| | | Cupreoboletus poikilochromus voucher MG 367a | Italy | KT157049 | KT157058 | | | |
| | | Cupreoboletus poikilochromus voucher TO HG 10091987 | Italy | KT157047 | KT157056 | | | |
| | | Cupreoboletus poikilochromus voucher TUR-A 208926 | Italy | MZ265177 | MZ265192 | | MZ277222 | MZ277233 |
| | | Cupreoboletus poikilochromus voucher TUR-A 208927 | Italy | MZ265178 | MZ265193 | | MZ277223 | |

| Genus | Species clade | Specimen | Country | ITS | LSU | RPB1 | RPB2 | TEF1 |
|---|---|---|---|---|---|---|---|---|
| | | Cyanoboletus poikilochromus voucher MB241005/01 | Italy | PV413925 | | | | |
| | | Cupreoboletus poikilochromus BdL8d17 | Portugal | | PZ127379 | | | |
| Cyanoboletus | abieticola | Cyanoboletus abieticola voucher Ayala-Vasquez (1010-ITVC) | Mexico | | MW750367 | | | |
| | | Cyanoboletus abieticola voucher Ayala-Vasquez (1128-ITVC) | Mexico | | MW750366 | | | |
| | | Cyanoboletus abieticola voucher Ayala-Vasquez (1129-ITVC) | Mexico | | MW750365 | | | |
| | | Cyanoboletus abieticola voucher Ayala-Vasquez (1141-ITVC) | Mexico | | MW750368 | | | |
| | | Cyanoboletus sp. OAV-2020a voucher AV33 | Mexico | | | | PP108649 | |
| | | Cyanoboletus sp. OAV-2020a voucher AV72 | Mexico | | | | PP108650 | |
| | | Cyanoboletus sp. OAV-2020a voucher Ferrusca390 | Mexico | MW750332 | MW750369 | | | |
| | | Cyanoboletus sp. OAV-2020a voucher MEXU-30106 | Mexico | MW209740 | | | | |
| | | Cyanoboletus sp. OAV-2020a voucher MEXU-30109 | Mexico | MW209738 | | | | |
| | | Cyanoboletus sp. OAV-2020a voucher MEXU-30111 | Mexico | MW209739 | | | | |
| | | Cyanoboletus sp. OAV-2020a voucher TB-2010-MEX 07 | Mexico | KC152077 | | | | |
| | | Cyanoboletus sp. OAV-2020a voucher TB-2010-MEX 08 | Mexico | KC152076 | | | | |
| | | Cyanoboletus sp. OAV-2020a voucher TB-2010-MEX 12 | Mexico | KC152075 | | | | |
| | brunneoruber | Boletus sp. HKAS 63504 | China | | KF112368 | KF112531 | KF112702 | KF112194 |
| | | Cyanoboletus brunneoruber voucher HKAS 52558 | China | KX904881 | KX904880 | KF739767 | KF739729 | KF739805 |
| | | Cyanoboletus brunneoruber voucher HKAS80579-1 | China | | KT990568 | KT990926 | KT990401 | KT990763 |
| | | Cyanoboletus brunneoruber voucher HKAS80579-2 | China | | KT990569 | KT990927 | KT990402 | KT990764 |
| | | Cyanoboletus brunneoruber voucher OR0233 | China | | | | MG212628 | MG212586 |
| | brunneoruber cf. | Cyanoboletus sp. voucher OR0491 | China | | | | MH614769 | MH614723 |
| | cyaneitinctus | Cyanoboletus cyaneitinctus voucher voucher B1915 | Canada | KY826057 | | | | |
| | | Cyanoboletus cyaneitinctus voucher CMMF002224 | Canada | KY826068 | | | | |
| | | Cyanoboletus cyaneitinctus voucher iNat66898219 | Canada | ON943338 | | | | |
| | | Boletus pulverulentus voucher JMP0012 | USA | EU819453 | | | | |
| | | Boletus pulverulentus voucher RT00004 | USA | EU819502 | | | | |
| | | Cyanoboletus aff. pulverulentes 'sp. IN01' voucher MICH139837 | USA | OM985877 | | | | |
| | | Cyanoboletus cyaneitinctus f. reticulatus voucher Farid 1035 | USA | MZ746113 | | | | |
| | | Cyanoboletus cyaneitinctus isolate OMDL bio-material iNAT:235571459 | USA | PX404192 | | | | |
| | | Cyanoboletus cyaneitinctus isolate OMDL iNat # 184805641 | USA | PP850729 | | | | |
| | | Cyanoboletus cyaneitinctus isolate OMDL MO # 551861 | USA | PX374970 | | | | |
| | | Cyanoboletus cyaneitinctus isolate OMDL MO # 552473 | USA | PX118628 | | | | |
| | | Cyanoboletus cyaneitinctus isolate S.D. Russell ONT Mushroom Observer 496468 | USA | OP749305 | | | | |
| | | Cyanoboletus cyaneitinctus voucher Farid 340 | USA | MW675739 | MW662574 | MW737502 | MW737461 | |

| Genus | Species clade | Specimen | Country | ITS | LSU | RPB1 | RPB2 | TEF1 |
|---|---|---|---|---|---|---|---|---|
| | | Cyanoboletus cyaneitinctus voucher Farid 920 | USA | MW675744 | MW662579 | MW737503 | MW737465 | |
| | | Cyanoboletus cyaneitinctus voucher JAB 184 | USA | MW675731 | MW662584 | MW737504 | MW737467 | |
| | | Cyanoboletus cyaneitinctus voucher JAB 324 | USA | MW675732 | MW662586 | MW737505 | MW737469 | |
| | | Cyanoboletus cyaneitinctus voucher JAB 325 | USA | MW675733 | | MW737506 | MW737470 | |
| | | Cyanoboletus cyaneitinctus voucher Mushroom Observer #323745 | USA | ON705304 | | | | |
| | | Cyanoboletus pulverulentus bio-material iNAT:55864593 | USA | MT939495 | | | | |
| | | Cyanoboletus pulverulentus voucher 9606 | USA | | KF030313 | KF030364 | | KF030418 |
| | | Cyanoboletus pulverulentus voucher iNat17028160 | USA | ON534188 | | | | |
| | | Cyanoboletus pulverulentus voucher iNat32191013 | USA | ON534167 | | | | |
| | | Cyanoboletus pulverulentus voucher MICH KUO-09220408 | USA | | MK601732 | | MK766294 | MK721086 |
| | | Cyanoboletus sp. strain B28, specimen voucher PRM 944518 | USA | LT714710 | MF373585 | | | |
| | | Uncultured Basidiomycota clone man24 soil H05 | USA | GU328532 | | | | |
| | | Uncultured ectomycorrhizal fungus isolate SR06T3-171 | USA | FM999526 | | | | |
| | hymenoglutinosus | Cyanoboletus hymenoglutinosus CAL DC 14-010 | India | NR_164239 | KT860060 | | | |
| | hymenoglutinosus sp. | Cyanoboletus sp. voucher OR0322 | Thailand | | | | MH614768 | MH614722 |
| | macroporus | Cyanoboletus sp. voucher DC 21-02 | India | OQ860238 | OQ860239 | | ON364552 | |
| | | Cyanoboletus sp. voucher DC 21-24 | India | OQ860240 | OQ860241 | | OQ876894 | |
| | | Cyanoboletus macroporus naseer | Pakistan | MW045557 | | | | |
| | | Cyanoboletus sp. isolate sarwar1 | Pakistan | MW369503 | | | | |
| | mediterraneensis | Cyanoboletus mediterraneensis voucher WC A 211022-06 | Greece | OR770584 | | | | |
| | | Cyanoboletus mediterraneensis K(M) 265123 | Israel | | NG_228932 | | | |
| | | Cyanoboletus mediterraneensis voucher K-M000265124 (ex herb. HAI A-001) | Israel | OM801199 | | | | |
| | | Cyanoboletus mediterraneensis soil TUE000513 | Italy | UDB01535438 | | | | |
| | | Cyanoboletus mediterraneensis soil TUE000523 | Italy | UDB01535455 | | | | |
| | | Cyanoboletus mediterraneensis soil TUF137111 | Italy | UDB07672757 | | | | |
| | | Cyanoboletus mediterraneensis voucher Hal-BP-25 | Italy | MT594497 | | | | |
| | | Cyanoboletus mediterraneensis voucher MS-1 | Italy | PP938742 | | | | |
| | | Cyanoboletus mediterraneensis voucher MS-2 | Italy | PP938743 | | | | |
| | | Cyanoboletus sp. PA-2021a voucher TUR-A 208928 | Italy | MZ265179 | MZ265194 | | MZ277224 | MZ277234 |
| | | Cyanoboletus sp. PA-2021a voucher TUR-A 208929 | Italy | MZ265181 | MZ265196 | | MZ277226 | MZ277236 |
| | | Cyanoboletus sp. PA-2021a voucher TUR-A 209100 | Italy | MZ265180 | MZ265195 | | MZ277225 | MZ277235 |
| | | Cyanoboletus sp. PA-2021a voucher TUR-A 209198 | Italy | MZ265182 | MZ265197 | | MZ277227 | MZ277237 |
| | | Cyanoboletus sp. PA-2021a voucher TUR-A 209199 | Italy | MZ265183 | MZ265198 | | MZ277228 | MZ277238 |
| | | Cyanoboletus mediterraneensis BdL3a04 | Portugal | PZ403734 | | | PZ164077 | PZ164082 |
| | | Cyanoboletus mediterraneensis M22#24 | Portugal | PZ157779 | | | | |

| Genus | Species clade | Specimen | Country | ITS | LSU | RPB1 | RPB2 | TEF1 |
|---|---|---|---|---|---|---|---|---|
| | | Cyanoboletus mediterraneensis M23#02 | Portugal | PZ395655 | PZ127384 | PZ164075 | PZ164081 | PZ164085 |
| | | Cyanoboletus mediterraneensis M25#18 | Portugal | PZ157780 | PZ127387 | | | |
| | | Cyanoboletus mediterraneensis M25#23 | Portugal | PZ157782 | PZ127386 | PZ164074 | PZ164078 | PZ164083 |
| | | Cyanoboletus mediterraneensis voucher VF171022CP1 | Portugal | PV170925 | | | | |
| | neotropicus | Boletus neotropicus voucher CFMR:BZ-3521 TJB-9821 | Belize | NR_175140 | NG_078673 | | | |
| | | Boletus sp. isolate AC689 | Panama | KY548897 | | | | |
| | | Boletus sp. isolate AC853 | Panama | KY548911 | | | | |
| | paurianus | Cyanoboletus paurianus voucher KD 22-008 | India | | OQ859920 | | OQ914389 | |
| | | Cyanoboletus paurianus voucher KD 22-009 | India | | NG_242126 | | OQ914388 | |
| | pulverulentus | Cyanoboletus pulverulentus WU-19504 | Austria | UDB0801853 | | | | |
| | | Cyanoboletus pulverulentus voucher RW109 | Belgium | | | | KT824013 | KT824046 |
| | | Boletus pulverulentus voucher A21 | China | JX434686 | | | | |
| | | Boletus pulverulentus voucher A7 | China | JX434685 | | | | |
| | | Cyanoboletus pulverulentus strain B23, specimen voucher PRM 944014 | Czech Republic | LT714705 | | | | |
| | | Cyanoboletus pulverulentus strain B24, specimen voucher PRM 944001 | Czech Republic | LT714706 | | | | |
| | | Cyanoboletus pulverulentus strain B25, specimen voucher PRM 944013 | Czech Republic | LT714707 | | | | |
| | | Cyanoboletus pulverulentus strain B26, specimen voucher PRM 944022 | Czech Republic | LT714708 | | | | |
| | | Cyanoboletus pulverulentus strain B27, specimen voucher PRM 935997 | Czech Republic | LT714709 | | | | |
| | | Cyanoboletus pulverulentus material sample BEECH PLOT27 | Germany | UDB0783407 | | | | |
| | | Cyanoboletus pulverulentus material sample BEECH PLOT9 | Italy | UDB0785546 | | | | |
| | | Cyanoboletus pulverulentus voucher 18188 | Italy | JF907794 | | | | |
| | | Cyanoboletus pulverulentus voucher MG 126a | Italy | KT157053 | KT157062 | | | |
| | | Cyanoboletus pulverulentus voucher MG 628a | Italy | KT157055 | KT157064 | | KT157069 | KT157073 |
| | | Cyanoboletus pulverulentus voucher TUR-A 208930 | Italy | MZ265186 | MZ265200 | | MZ277230 | |
| | | Cyanoboletus pulverulentus voucher TUR-A 208931 | Italy | MZ265185 | MZ265199 | | MZ277229 | |
| | | Boletus pulverulentus voucher CA050916-04 | Italy | HM347646 | | | | |
| | | Cyanoboletus pulverulentus specimen O-F-251505 | Norway | UDB036254 | | | | |
| | | Cyanoboletus pulverulentus strain B21, specimen voucher PRM 935923 | Portugal | LT714704 | | | | |
| | | Cyanoboletus pulverulentus voucher MG 456a | Portugal | KT157054 | KT157063 | | | |
| | | Cyanoboletus pulverulentus voucher VF270922SS1 | Portugal | PV461258 | | | | |
| | | Cyanoboletus pulverulentus voucher LUGO:ECC18060709 | Spain | MW376644 | | | | |
| | | Uncultured Cyanoboletus clone 12944 | Spain | MW282397 | | | | |
| | | Uncultured Cyanoboletus clone 13143 | Spain | MW282441 | | | | |

| Genus | Species clade | Specimen | Country | ITS | LSU | RPB1 | RPB2 | TEF1 |
|---|---|---|---|---|---|---|---|---|
| | | Uncultured Cyanoboletus clone 14181 | Spain | MW282401 | | | | |
| | | Uncultured Cyanoboletus clone 15892 | Spain | MW282492 | | | | |
| | | Cyanoboletus pulverulentus strain U3750 | Sweden | PP455303 | | | | |
| | | Cyanoboletus pulverulentus specimen AH97030 | UK | UDB000408 | | | | |
| | | Cyanoboletus pulverulentus voucher HFRG PC241026 2 | UK | PV690293 | | | | |
| | pulverulentus aff. | Cyanoboletus pulverulentus TN1601 | Japan | LC832002 | | | | |
| | | Boletus pulverulentus voucher ASIS22672 | South Korea | KP004920 | | | | |
| | sinopulverulentus | Boletus sinopulverulentus voucher HMAS 266894 holotype | China | KC579402 | | | | |
| | | Cyanoboletus sp. NM-2023a voucher BJTC FM2319-A holotype | China | NR_191306 | NG_243401 | | OR659937 | OR659986 |
| | | Cyanoboletus sp. NM-2023a voucher BJTC FM2319-B | China | | OR655226 | | OR659976 | OR660025 |
| | | Cyanoboletus sp. HKAS 90208-1 | China | | KT990571 | | KT990404 | KT990766 |
| | | Cyanoboletus sp. HKAS 90208-2 | China | | | | KT990405 | KT990767 |
| | sinopulverulentus aff. (1) | Boletus sp. HKAS 52639 | China | | KF112367 | KF112530 | KF112701 | KF112195 |
| | | Cyanoboletus sp. HKAS 59418 | China | | KT990570 | | KT990403 | KT990765 |
| | | Cyanoboletus sp. voucher OR0257 | China | | | | MG212629 | MG212587 |
| | | Cyanoboletus sp. TN201707 | Japan | LC832054 | | | | |
| | | Cyanoboletus flavocontextus voucher LE F-344051 | Viet Nam | PP317924 | PP313111 | | | PP320320 |
| | | Cyanoboletus flavocontextus voucher LE F-344052 | Viet Nam | PP317923 | | | | PP320319 |
| | sinopulverulentus aff. (2) | Boletus sp. HKAS 59609 | China | | KF112366 | KF112529 | KF112700 | KF112193 |
| | | Cyanoboletus sinopulverulentus DC 16-51 | India | | MH684757 | | | |
| | | Cyanoboletus sp. voucher OR0961 | Thailand | | | | MH614770 | MH614724 |
| | viscidiceps | Cyanoboletus viscidiceps HMJAU68168 (R345) | China | | OR673995 | OR683473 | OR683479 | OR683468 |
| | sp. sulcatipes | Cyanoboletus sp. sulcatipes 47 1 | Pakistan | PX783209 | | | | |
| | | Cyanoboletus sp. sulcatipes 47 2 | Pakistan | PX783210 | | | | |
| | sp. | Boletus sp. KA12-1240 | South Korea | KR673493 | | | | |

### Table S2. List of Genbank accession numbers for the outgroup sequences

| Genus | Specimen | ITS | LSU | RPB1 | RPB2 | TEF1 |
|---|---|---|---|---|---|---|
| Boletus s.l. | Boletus roseoareolatus voucher CFMR:BZ-3554 TJB-9854 | MN250202 | MN250177 | | | |
| | Boletus rugulosiceps voucher CFMR:BZ-3533 TJB-9833 | | MN250178 | | | |
| | Boletus sp. voucher JD0693 Burundi | | | | MH645599 | MH645591 |
| | uncultured Boletaceae Guinea | AM113453 | | | | |
| Amoenoboletus | Amoenoboletus granulopunctatus voucher HKAS80250 | | MW520185 | | MW560080 | MW566746 |
| Baorangia | Baorangia duplicatopora voucher N.K.Zeng4511 (FHMU5879) | | MW473484 | | | MW485969 |
| | Baorangia pseudocalopus voucher HKAS 75739 | | KJ184558 | KJ184564 | KM605179 | KJ184570 |
| | Baorangia rufomaculata voucher 4414 | | KF030248 | KF030369 | | KF030406 |
| Butyriboletus | Butyriboletus appendiculatus strain Bap1 | | AF456837 | KF030359 | | JQ327025 |
| | Butyriboletus roseoflavus voucher HKAS:54099 | | KF739665 | KF739741 | KF739703 | KF739779 |
| Caloboletus | Caloboletus aff. calopus HKAS 74739 | | KF112335 | KF112507 | KF112667 | KF112166 |
| | Caloboletus calopus voucher BR5020159063805 | | KJ184554 | KJ184560 | | KJ184566 |
| | Caloboletus firmus voucher CFMR BZ-1721 | | MK601726 | | MK766288 | MK721080 |
| | Caloboletus griseoflavus voucher BJTC FM699 holotype | | OR655181 | | OR659933 | OR659982 |
| | Caloboletus guanyui voucher N.K.Zeng3257 (FHMU2218) | | MH879705 | | MH879748 | MH879732 |
| | Caloboletus inedulis voucher MICH KUO-07031403 | | MK601727 | | MK766289 | MK721081 |
| | Caloboletus panniformis strain HKAS77506 | | KJ605669 | KJ619477 | | KJ619468 |
| | Caloboletus radicans voucher HKAS80856 | | KJ184557 | KJ184563 | | KJ184569 |
| | Caloboletus xiangtoushanensis voucher GDGM44833 | | KY800415 | KY800425 | | KY800418 |
| | Caloboletus yunnanensis voucher Hao230 HKAS69214 | | KJ184556 | KJ184562 | KT990396 | KJ184568 |
| Costatisporus | Costatisporus cyanescens Henkel 9061 | | | LC053663 | LC053664 | |
| | Costatisporus cyanescens Henkel 9067 | | LC053662 | | | |
| Crocinoboletus | Crocinoboletus rufoaureus voucher HKAS:53424 | | KF112435 | | KF112710 | KF112206 |
| Erythrophylloporus | Erythrophylloporus cinnabarinus voucher GDGM46541 | | MH374043 | MH374030 | MH374034 | |
| Exsudoporus | Exsudoporus frostii voucher NY815462 | | JQ924342 | | KF112675 | KF112164 |
| | Exsudoporus ruber voucher KUN-HKAS 106891 (Zhu L. Yang6279) | | MN930518 | MT063118 | MT063120 | MT063123 |
| Gastroboletus | Gastroboletus thibetanus HKAS 57093 | | KF112326 | KF112496 | KF112655 | |
| Gymnogaster | Gymnogaster boletoides voucher NY01194009 | | KT990572 | KT990928 | KT990406 | KT990768 |
| Hongoboletus | Hongoboletus ventricosus voucher HKAS:63598 | | KF112317 | KF112502 | KF112663 | KF112152 |
| Imperator | Imperator torosus strain ZK193 | | MW521084 | | | |
| | Imperator torosus voucher MB000258 | | | | MW560082 | MW566748 |
| Lanmaoa | Boletaceae sp voucher CNV21 | MT345198 | | | | |
| | Boletus fragrans MA-Fungi 477 | AJ419186 | | | | |

| Genus | Specimen | ITS | LSU | RPB1 | RPB2 | TEF1 |
|---|---|---|---|---|---|---|
| (Lanmaoa) | Boletus sensibilis FLAS-F-61203 | MH211808 | | | | |
| | Boletus sensibilis var. subviscidus voucher 3929 | | KF030310 | | | |
| | Boletus sensibilis voucher CMMF002483 | KY826075 | | | | |
| | Boletus sensibilis voucher iNAT 15950523 | ON166648 | | | | |
| | Boletus sensibilis voucher MICH10045 | PX898852 | | | | |
| | Boletus sensibilis voucher Mushroom Observer 288555 | MH257571 | | | | |
| | Boletus sensibilis voucher Mushroom Observer 424678 | ON950102 | | | | |
| | Boletus sp MGW1536 Tennessee USA | MG663300 | | | | |
| | Boletus sp. HKAS 55445 China | | KF739672 | KF739748 | KF739710 | KF739786 |
| | Boletus sp. MN-2013a voucher DPL6899 | | KF030260 | | | |
| | Boletus sp. strain MGW1536 | | MF797695 | | | |
| | Butyriboletus appendiculatus LEB Fungi-1597 | FM958176 | | | | |
| | Lanmaoa angustispora voucher HKAS 74752 | | KM605139 | KM605166 | KM605177 | KM605154 |
| | Lanmaoa angustispora voucher SWAT003369 | PQ870408 | | | | |
| | Lanmaoa asiatica HKAS 54094 | NR_201094 | KF112353 | KF112522 | KF112682 | KF112161 |
| | Lanmaoa carminipes voucher 4591 | | KF030259 | | | |
| | Lanmaoa carminipes voucher MB06-001 | | JQ327001 | | | |
| | Lanmaoa carminipes voucher MICH10002 | PX898847 | | | | |
| | Lanmaoa cf borealis voucher AB35 | MH796994 | | | | |
| | Lanmaoa flavorubra voucher NY775777 | | JQ924339 | | KF112681 | KF112160 |
| | Lanmaoa fragrans AT2000183 | UDB000946 | | | | |
|  | Lanmaoa fragrans isolate sporocarp Coimbra | MN314115 |  |  |  |  |
| | Lanmaoa fragrans M24#34 | PZ403735 | | | | |
| | Lanmaoa fragrans voucher 18555 | JF907800 | | | | |
| | Lanmaoa fragrans voucher TO HG061002 | | MH036176 | | | |
| | Lanmaoa pallidorosea voucher DUKE 0351606 | OL342405 | | | | |
| | Lanmaoa roseocrispans voucher ARB92-10 | | KP327615 | | | |
| | Lanmaoa sp bio-material MUOB 370068 | OL142184 | | | | |
| | Lanmaoa sp fallax isolate SD Russell ONT iNaturalist 127287218 | OP549259 | | | | |
| | Lanmaoa sp fallax voucher Mushroom Observer 333824 | ON705295 | | | | |
| | Lanmaoa sp isolate YAAS40LA | OR075198 | | | | |
| | Lanmaoa sp voucher MES-3758 | ON383391 | | | | |
| | Lanmaoa sp voucher Mushroom Observer 244162 | MH257551 | | | | |
| | Lanmaoa sp voucher Mushroom Observer 327563 (MycoFlora 54402) | MW882066 | | | | |
| | Lanmaoa sublurida isolate SD Russell ONT iNaturalist #129868866 | OP470595 | | | | |

| Genus | Specimen | ITS | LSU | RPB1 | RPB2 | TEF1 |
|---|---|---|---|---|---|---|
| | Lanmaoa sublurida voucher Farid 1023 | | MW662572 | | | |
| Neoboletus | Neoboletus brunneorubrocarpus voucher HKAS126559 | | OQ888720 | OQ873475 | OQ873496 | OQ873457 |
| | Neoboletus ferrugineus voucher HKAS77617 | | KT990595 | KT990943 | KT990430 | KT990788 |
| | Neoboletus flavosanguineus voucher TUR-A 208905 | | MZ265201 | | MZ277231 | MZ277239 |
| | Neoboletus hainanensis voucher HKAS74880 | | KT990597 | KT990945 | KT990432 | KT990790 |
| | Neoboletus obscureumbrinus voucher HKAS63498 | | KT990598 | KT990946 | KT990433 | KT990791 |
| Pulchroboletus | Boletus neoregius isolate NY796166 | JQ924304 | JQ924344 | | | |
| | Pulchroboletus pseudosclerotiorum voucher Mushroom Observer #215764 | | MH220316 | | | |
| | Pulchroboletus roseoalbidus voucher MCVE 18217 | KJ729488 | | | | |
| | Pulchroboletus rubricitrinus Farid335 | MF193884 | ENA\|MG026638 | ENA\|UDM69794 | ENA\|UDM69748 | |
| | Pulchroboletus rubricitrinus isolate SD Russell iNat#197547868 | PV223691 | | | | |
| | Pulchroboletus sclerotiorum isolate SD Russell ONT iNaturalist #129632763 | OP749126 | | | | |
| | Pulchroboletus sclerotiorum voucher FLAS-F-60333 | MF098659 | MF614166 | MF614168 | MF614169 | MF614167 |
| Pulveroboletus | Pulveroboletus macrosporus voucher HKAS59530 | | KT990617 | KT990967 | KT990452 | KT990811 |
| Rubroboletus | Rubroboletus dupainii voucher JAM 0607 | | KF030251 | KF030361 | | KF030413 |
| | Rubroboletus esculentus strain K.Zhao893 | | KY272129 | KY272132 | KY272135 | KY272138 |
| Rugiboletus | Rugiboletus aff. extremiorientale HKAS 68586 | | KF112402 | KF112534 | KF112719 | KF112197 |
| Singerocomus | Singerocomus inundabilis Aime 4004 | | LC043090 | LC043091 | LC043092 | |
| Suillellus | Suillellus aff. amygdalinus HKAS 57262 | | KF112316 | KF112501 | KF112660 | KF112174 |
| | Suillellus amygdalinus voucher NY00035656 | | KT990650 | KT990990 | KT990477 | KT990840 |
| | Suillellus flaviporus strain HKAS126551 | | OQ888725 | OQ873480 | OQ873500 | OQ873462 |
| | Suillellus sp. NM-2023a voucher BJTC FM1755 holotype | | OR655212 | | OR659964 | OR660011 |
| Sutorius | Sutorius aff. eximius HKAS 52672 | | KF112399 | KF112584 | KF112802 | KF112207 |
| | Sutorius brunneissimus voucher HKAS:52660 | | KF112314 | KF112492 | KF112650 | KF112143 |
| | Sutorius eximius voucher HKAS59657 | | KT990707 | KT991029 | KT990505 | KT990887 |
| | Sutorius magnificus voucher HKAS:74939 | | KF112320 | KF112494 | KF112653 | KF112148 |
| Tropicoboletus | Boletus ruborculus voucher JBSD133072-ANGE203 | OQ108296 | | | | |
| | Boletus ruborculus voucher NY 577594-TJB 8253 (Holotype) | OQ108299 | | | | |
| | Pulchroboletus sp soil Cuba | KIGB01028760 | | | | |
| | Tropicoboletus ruborculus voucher JBSD133074 (ANGE1406) | OQ108297 | OQ102359 | | OQ117431 | OQ110624 |
| | Tropicoboletus ruborculus voucher MO439745-komille277 | OQ108298 | OQ102361 | | OQ117432 | OQ110625 |

#### Table S3. List of Genbank accession numbers for the *Neoboletus* sequences in Figure S7

| Specimen | ITS | LSU | RPB2 | TEF1 |
|---|---|---|---|---|
| Boletaceae sp. voucher SD Russell MycoMap 6640 | MK560094 | | | |
| Boletus cf. subvelutipes voucher Mushroom Observer #206608 | MH220323 | MH220333 | | MH318609 |
| Boletus cf. subvelutipes voucher Mushroom Observer #212359 | | MH236217 | | MH337288 |
| Boletus flavosanguineus voucher TUR-A 208905 | MZ265187 | MZ265201 | MZ277231 | MZ277239 |
| Boletus roseobadius voucher MICH10032 holotype | PZ253648 | | | |
| Boletus rufocinnamomeus voucher MICH10042 holotype | PZ253655 | | | |
| Boletus sp. HKAS 53719 | | KF739677 | KF739715 | KF739791 |
| Boletus sp. HKAS 55455 | | KF739678 | KF739716 | KF739792 |
| Boletus sp. HKAS 59443 | | KF739684 | KF739722 | KF739798 |
| Boletus sp. HKAS 74728 | | KF739676 | KF739714 | KF739790 |
| Boletus subluridellus isolate 3693 | KM248927 | | | |
| Boletus subluridellus isolate OMDL iNat #178099705 | PQ822133 | | | |
| Boletus subluridellus isolate OMDL iNat #178605534 | PP850490 | | | |
| Boletus subluridellus isolate OMDL iNat #179300453 | PQ847559 | | | |
| Boletus subluridellus isolate OMDL iNat #179825466 | PQ822103 | | | |
| Boletus subluridellus isolate OMDL iNat #181939192 | PQ287891 | | | |
| Boletus subvelutipes voucher NYSf3093 | ON142311 | ON142311 | OQ680486 | |
| Boletus vermiculosoides voucher HRL0837 | KY826167 | | | |
| Gastroboletus citrinobrunneus voucher NY65239 | PZ258852 | | | |
| Gastroboletus sp. HKAS 57093 | | KF112326 | KF112655 | |
| Gastroboletus sp. JLF2137 | JX415334 | | | |
| Gastroboletus sp. JLF3420 | KU160177 | | | |
| Gastroboletus sp. voucher JLF4412 | | MH216147 | | |
| Gastroboletus turbinatus | | AF336248 | | |
| Gastroboletus turbinatus isolate CA FUNDIS iNaturalist #161348882 | OR778476 | | | |
| Gastroboletus turbinatus var flammeus isolate CA FUNDIS iNaturalist #170577389 | OR858671 | | | |
| Gastroboletus turbinatus var flammeus isolate iNaturalist #174956144 | PP659827 | | | |
| Gastroboletus turbinatus var flammeus voucher MICH5547 holotype | PZ253745 | | | |
| Gastroboletus vividus voucher JLF4456 | MH213058 | | | |
| Gastroboletus vividus voucher JLF8924 CSNM iNaturalist #58926871 | PZ287843 | PZ287844 | | |
| Neoboletus erythropus voucher FBozok00135 | MK272852 | | | |
| Neoboletus sp. 'chaemeleon' isolate OMDL iNat #232931767 | PX369697 | | | |
| Neoboletus sp. 'flammeus' isolate CA FUNDIS iNaturalist #230651893 | PV587952 | | | |
| Neoboletus sp. voucher Arora21771 | PV055918 | | | |
| Neoboletus sp. voucher Arora21773b | PV055922 | | | |

| Specimen | ITS | LSU | RPB2 | TEF1 |
|---|---|---|---|---|
| Neoboletus sp. voucher Arora21786 | PV055913 | | | |
| Neoboletus sp. voucher JLF9996 iNaturalist #105935935 | PV052870 | | | |
| Neoboletus angiocarpus voucher KD 24HP-134 type | | PQ578768 | PQ613843 | PQ613840 |
| Neoboletus angiocarpus voucher KD 24HP-142 | | PQ578864 | PQ613842 | PQ613841 |
| Neoboletus sp. voucher Mushroom Observer #287453 | MK533786 | MK537316 | | |
| Neoboletus sp. voucher Mushroom Observer #251141 | | MH234861 | | MH337280 |
| Neoboletus sp. voucher NS4910 | PV055948 | | | |
| Neoboletus subvelutipes voucher Mushroom Observer #285181 | MH244205 | MH244204 | | MH347318 |

**Fig. S1 nucLSU +RPB1 +RPB2 +TEF1**

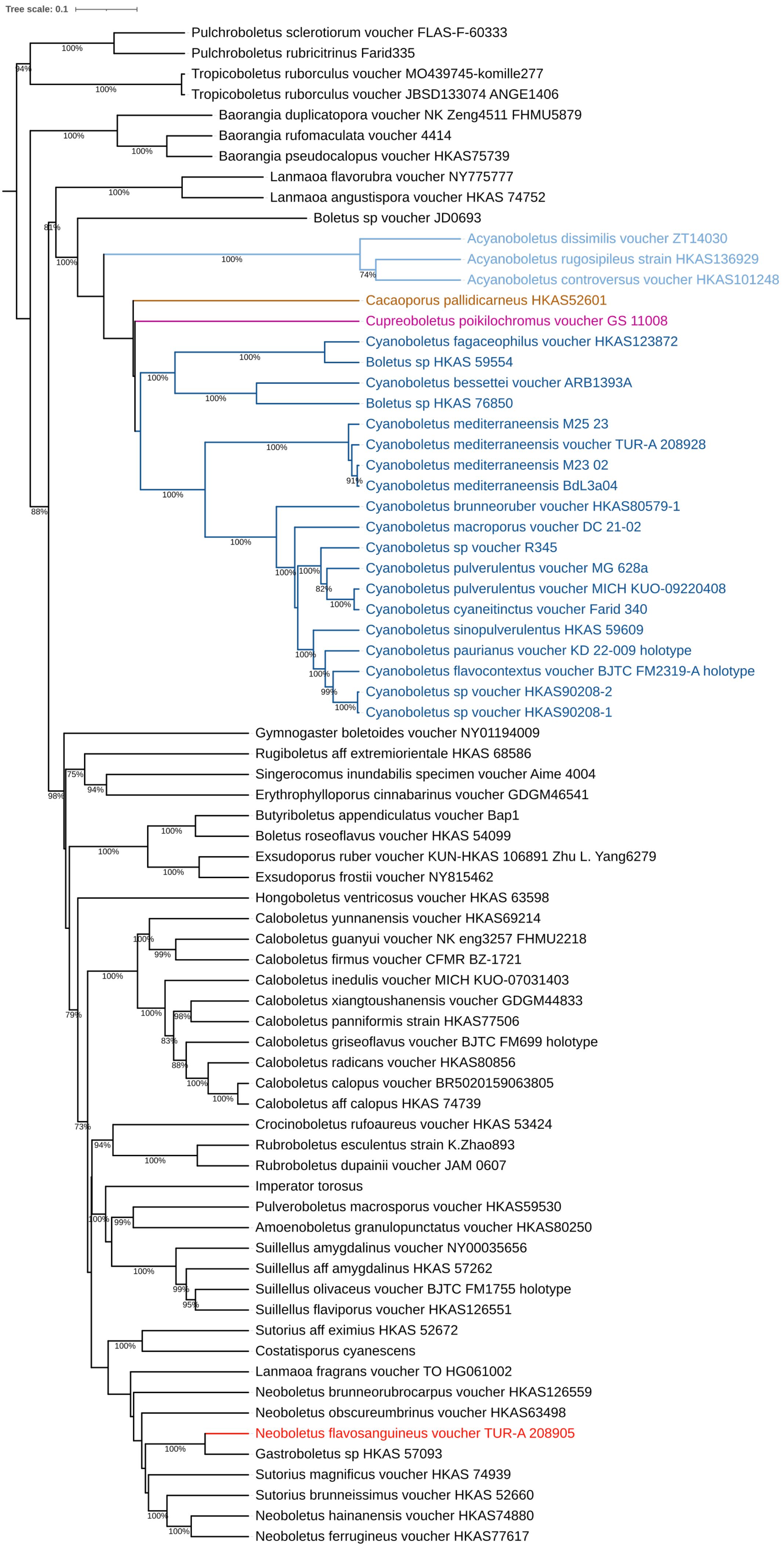

### Fig. S2
### a) nucLSU

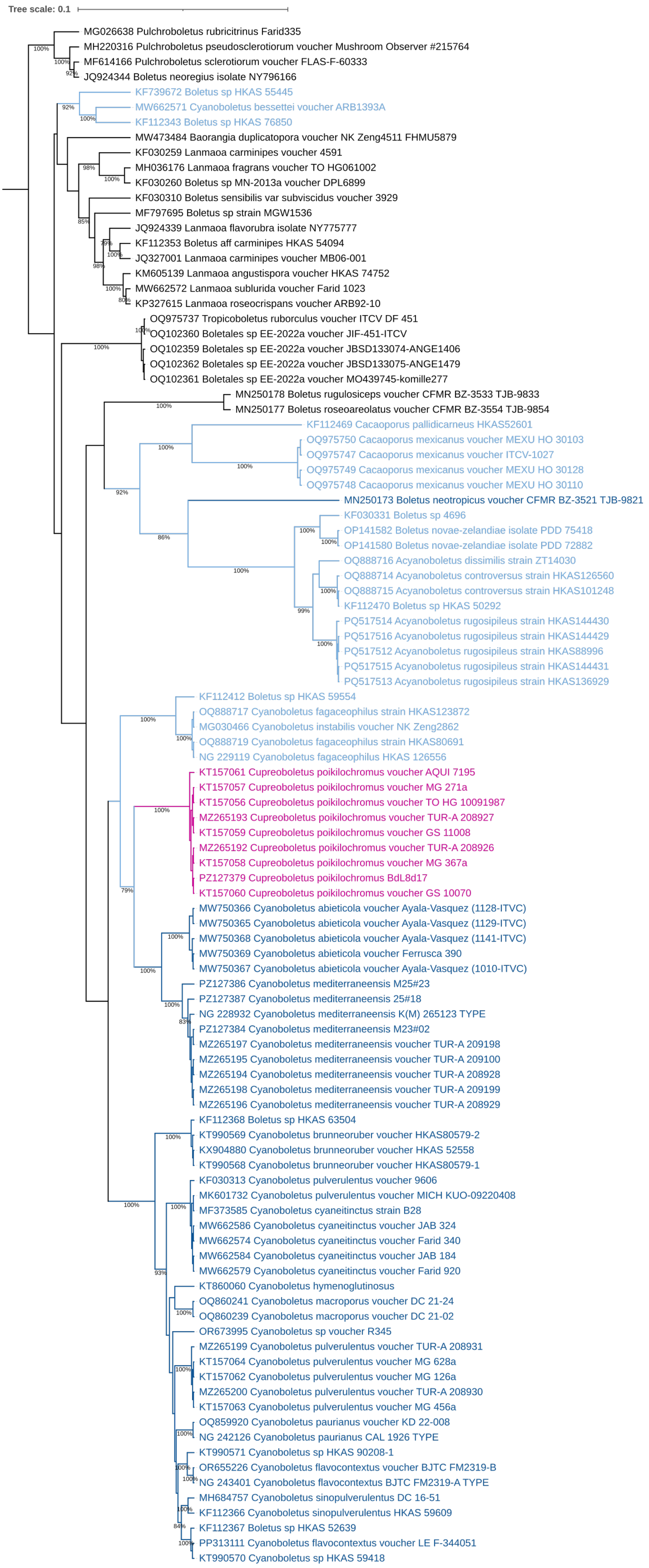

#### Fig. S2 (continued)

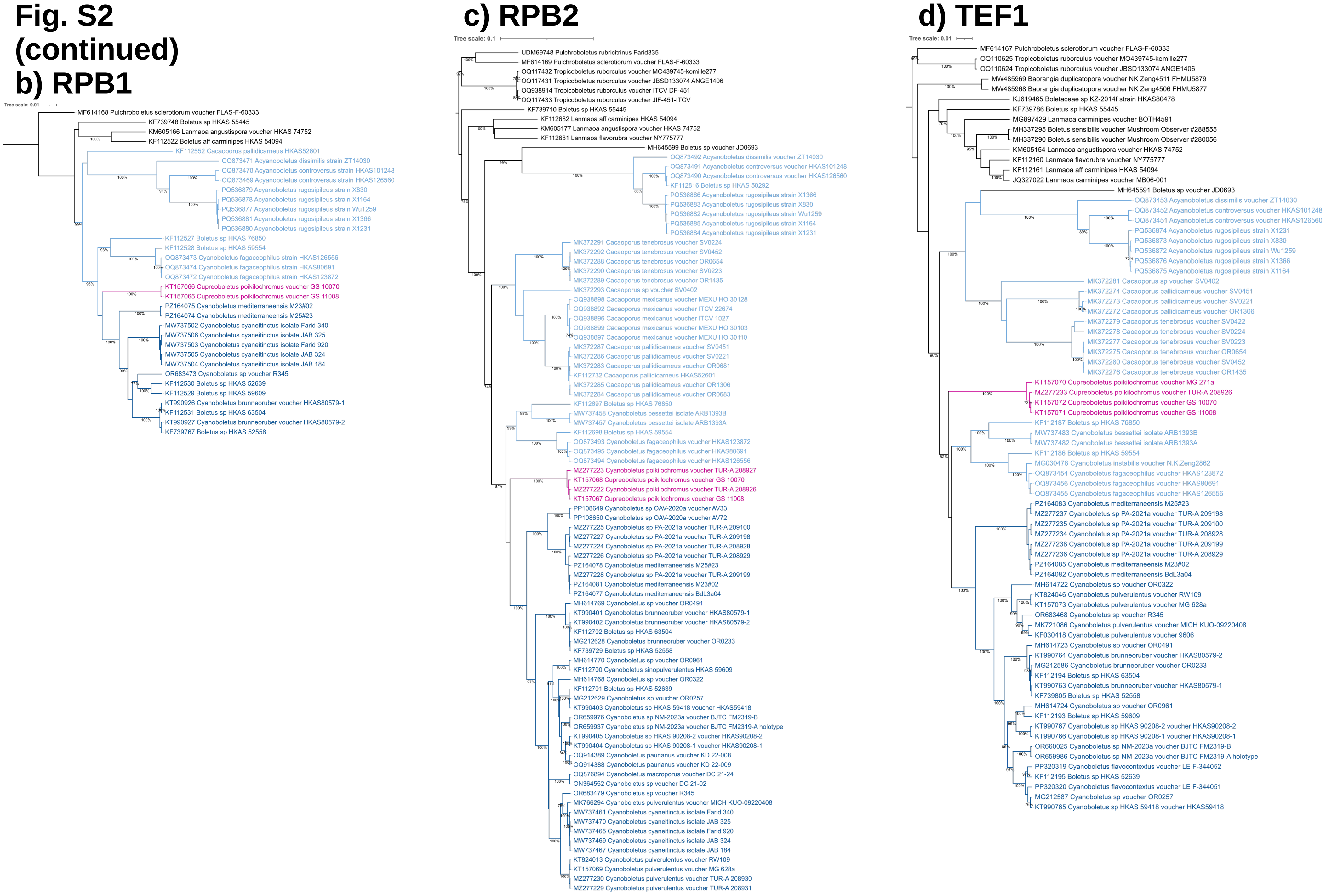

### Fig. S3 ITS+ nucLSU

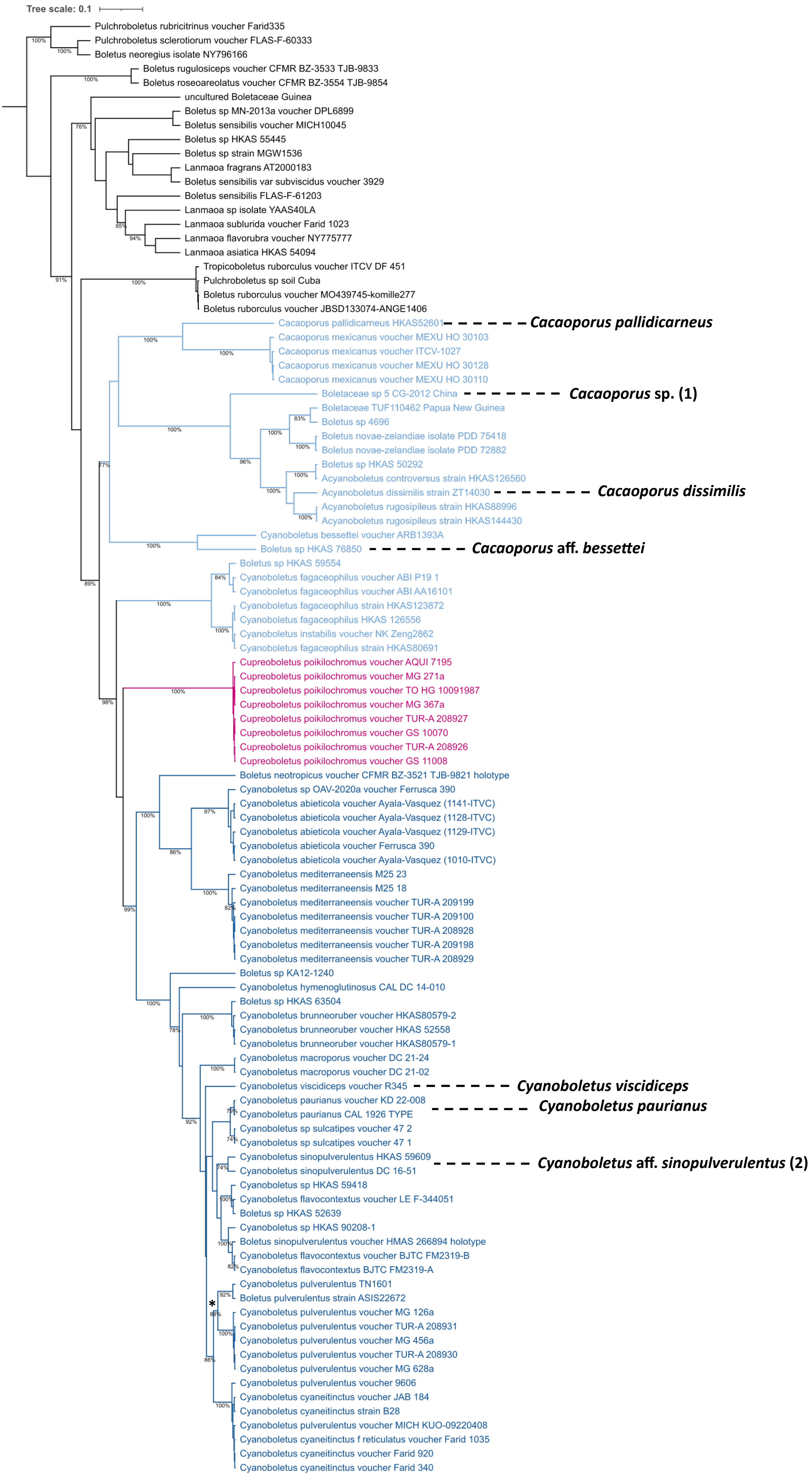

**Fig. S4**
**nucLSU**
**+RPB2**
**+TEF1**

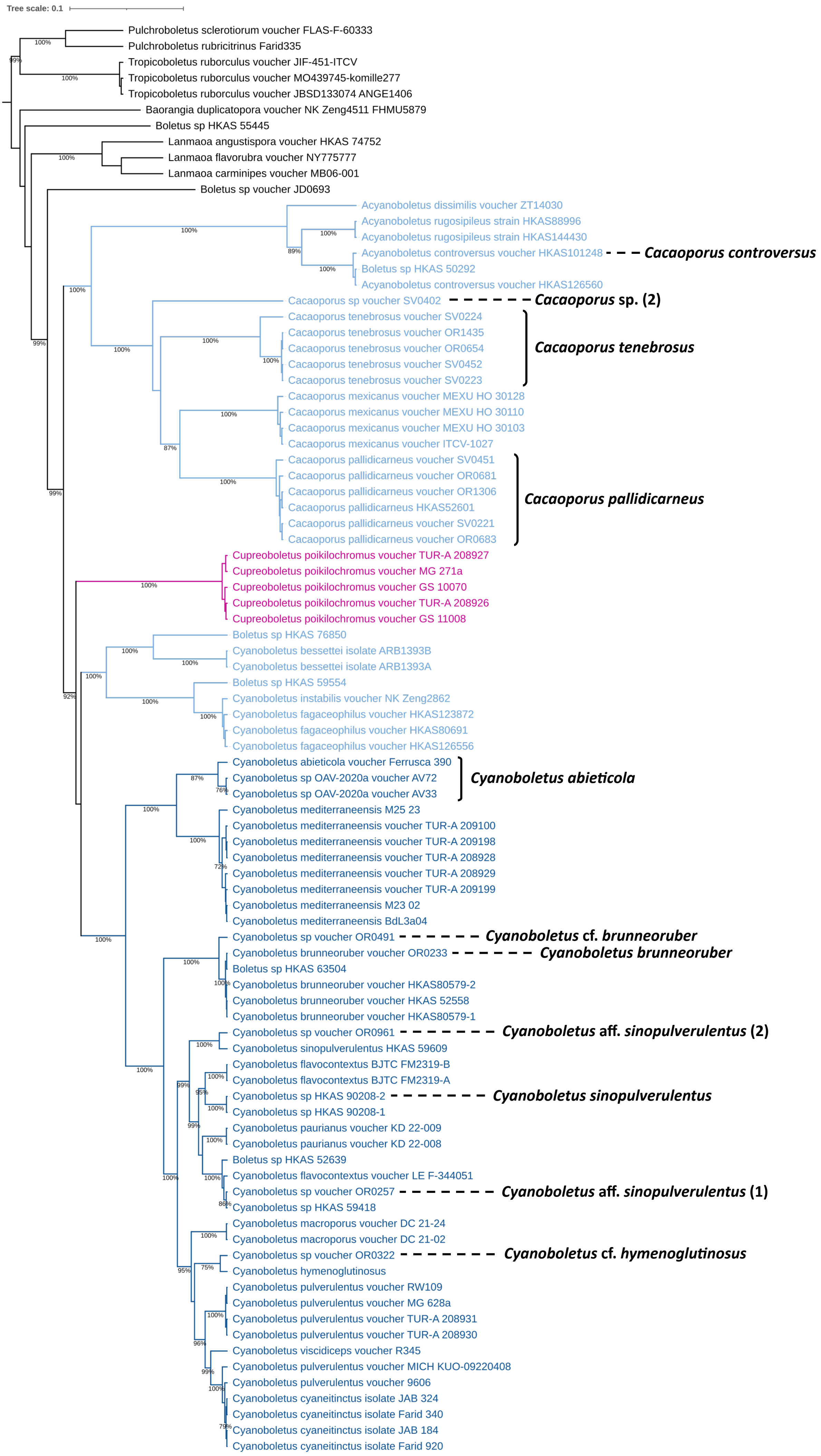

**Fig. S5**
**ITS**

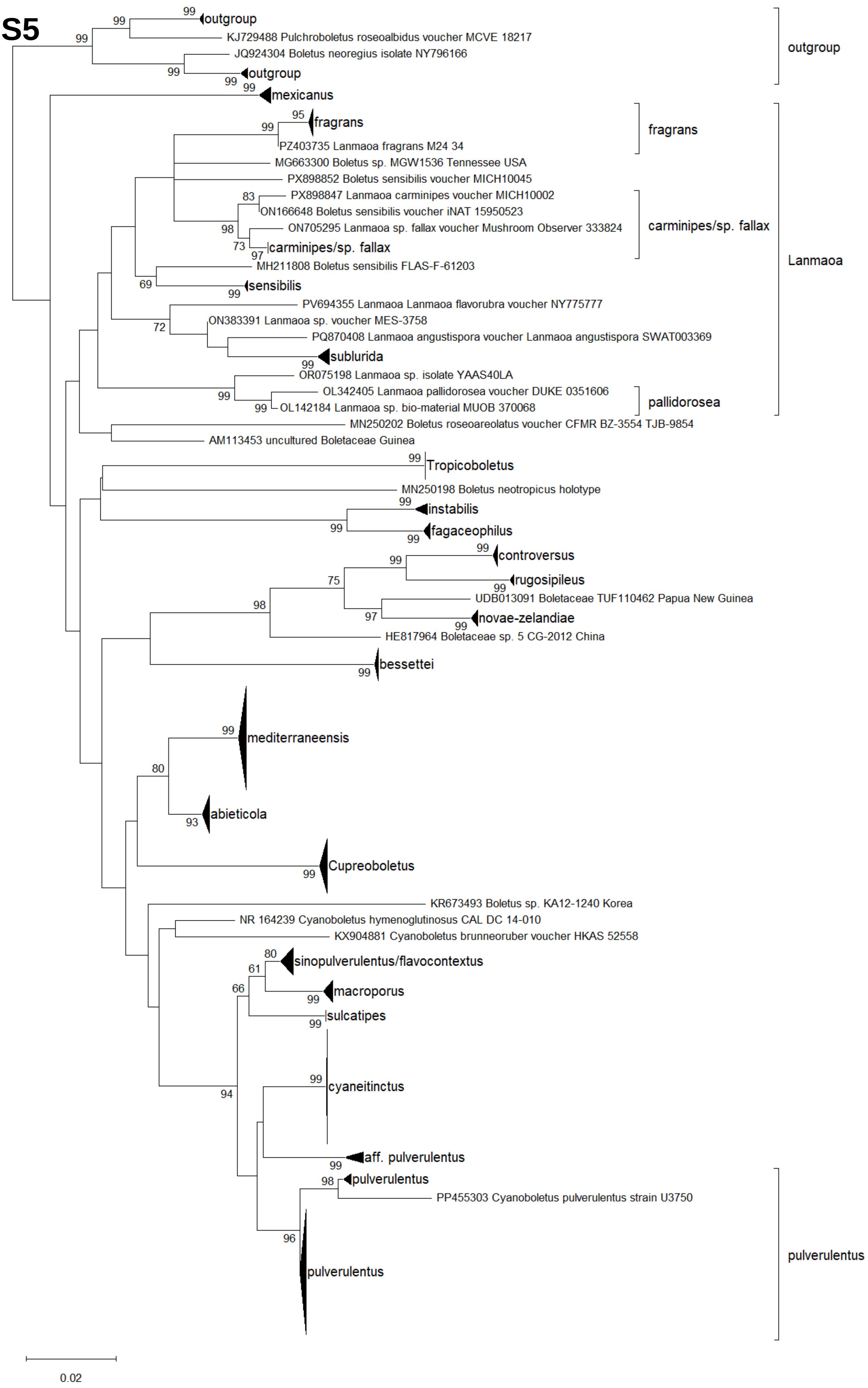
outgroup
KJ729488 Pulchroboletus roseoalbidus voucher MCVE 18217
JQ924304 Boletus neoregius isolate NY796166
outgroup
mexicanus
fragrans
PZ403735 Lanmaoa fragrans M24 34
MG663300 Boletus sp. MGW1536 Tennessee USA
PX898852 Boletus sensibilis voucher MICH10045
PX898847 Lanmaoa carminipes voucher MICH10002
ON166648 Boletus sensibilis voucher iNAT 15950523
ON705295 Lanmaoa sp. fallax voucher Mushroom Observer 333824
carminipes/sp. fallax
MH211808 Boletus sensibilis FLAS-F-61203
sensibilis
PV694355 Lanmaoa Lanmaoa flavorubra voucher NY775777
ON383391 Lanmaoa sp. voucher MES-3758
PQ870408 Lanmaoa angustispora voucher Lanmaoa angustispora SWAT003369
sublurida
OR075198 Lanmaoa sp. isolate YAAS40LA
OL342405 Lanmaoa pallidorosea voucher DUKE 0351606
OL142184 Lanmaoa sp. bio-material MUOB 370068
MN250202 Boletus roseoareolatus voucher CFMR BZ-3554 TJB-9854
AM113453 uncultured Boletaceae Guinea
Tropicoboletus
MN250198 Boletus neotropicus holotype
instabilis
fagaceophilus
controversus
rugosipileus
UDB013091 Boletaceae TUF110462 Papua New Guinea
novae-zelandiae
HE817964 Boletaceae sp. 5 CG-2012 China
bessettei
mediterraneensis
abieticola
Cupreoboletus
KR673493 Boletus sp. KA12-1240 Korea
NR 164239 Cyanoboletus hymenoglutinosus CAL DC 14-010
KX904881 Cyanoboletus brunneoruber voucher HKAS 52558
sinopulverulentus/flavocontextus
macroporus
sulcatipes
cyaneitinctus
aff. pulverulentus
pulverulentus
PP455303 Cyanoboletus pulverulentus strain U3750
pulverulentus
outgroup
fragrans
carminipes/sp. fallax
Lanmaoa
pallidorosea
pulverulentus
0.02

### Online Resource 7

#### Table S4. Correspondence between unidentified clades of *Cyanoboletus* and *Cacaoporus* in Biketova et al. (2026) and the present work (Fig. 1, Fig. S3/Online Resource 4, Table S1/Online Resource 1).

| Clade | Present work | Specimens in present work |
|---|---|---|
| sp. 1 | *Cyanoboletus* aff. *pulverulentus* | TN1601 Japan, ASIS22672 |
| sp. 2 | *Cyanoboletus* cf. *hymenoglutinosus* | OR0322 |
| sp. 3 | *Cyanoboletus* aff. *sinopulverulentus* (1) | HKAS 52639, HKAS 59418, OR0257, LE F-344051, LE F-344052 |
| sp. 4 | *Cyanoboletus sinopulverulentus* | HKAS 90208-1, HKAS 90208-2 |
| sp. 5 | *Cyanoboletus* aff. *sinopulverulentus* (2) | DC 16-51, OR0961 |
| sp. 6 | *Cyanoboletus* cf. *brunneoruber* | OR0491 |
| sp. 7 | *Cacaoporus* aff. *bessettei* | HKAS 76850 |

### Fig. S6

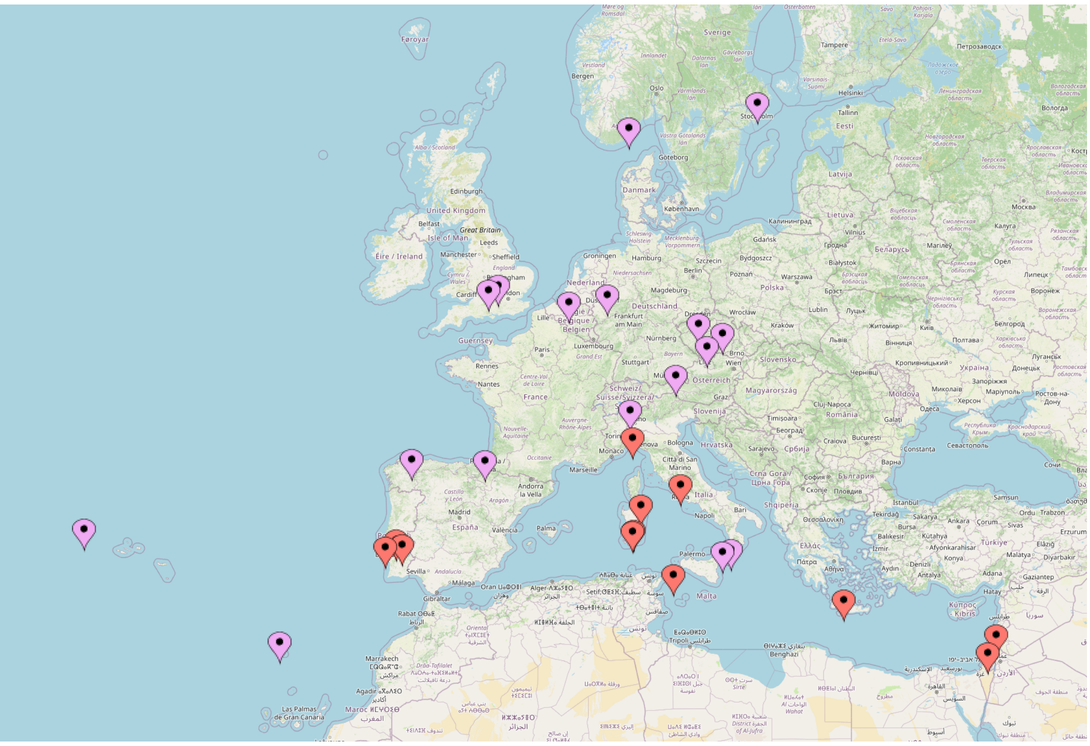

*Cyanoboletus pulverulentus* (violet dots) and *Cyanoboletus mediterraneensis* (red dots).

### Fig. S7 a)
### ITS

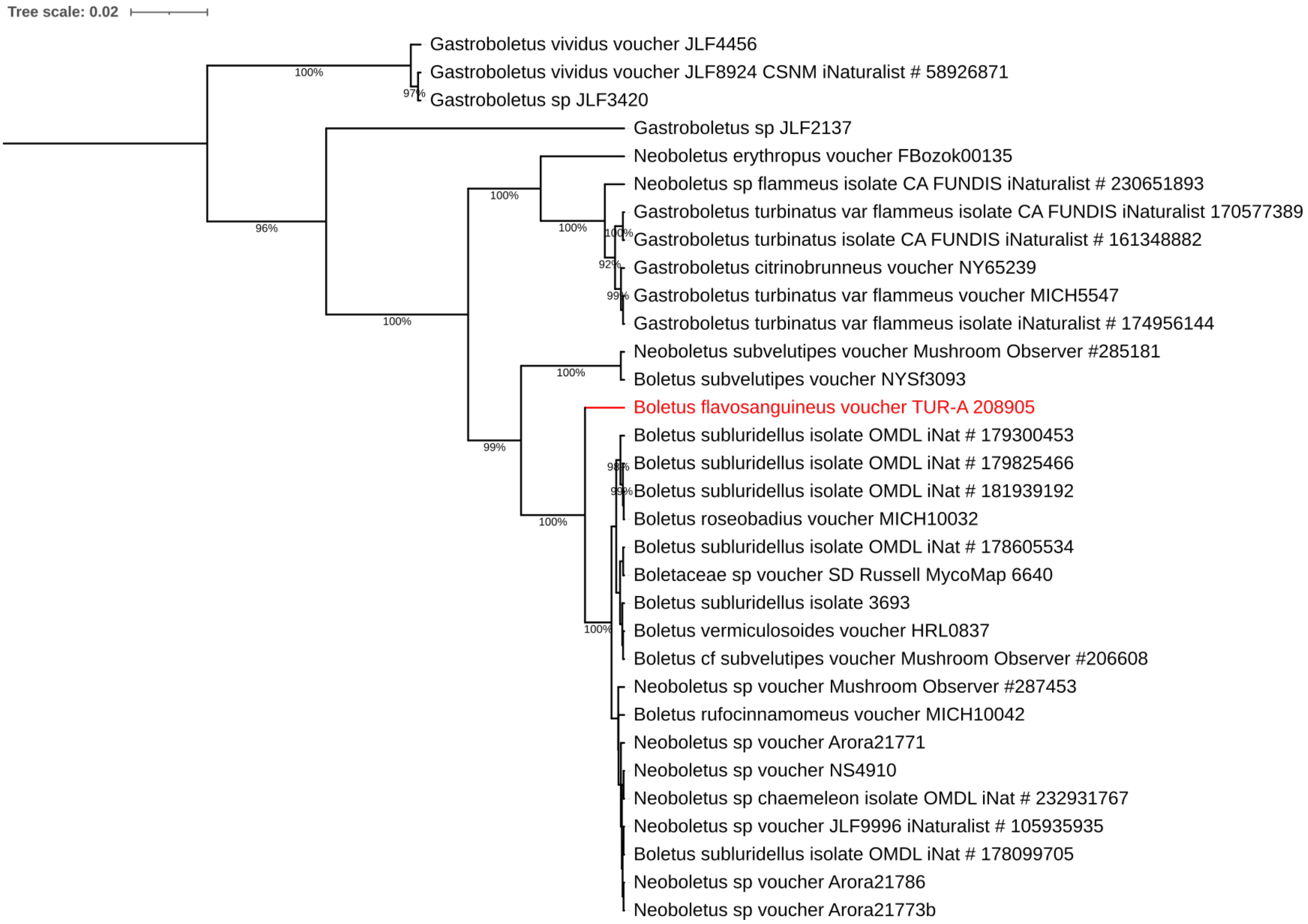


# b)
### nucLSU+RPB2+TEF1

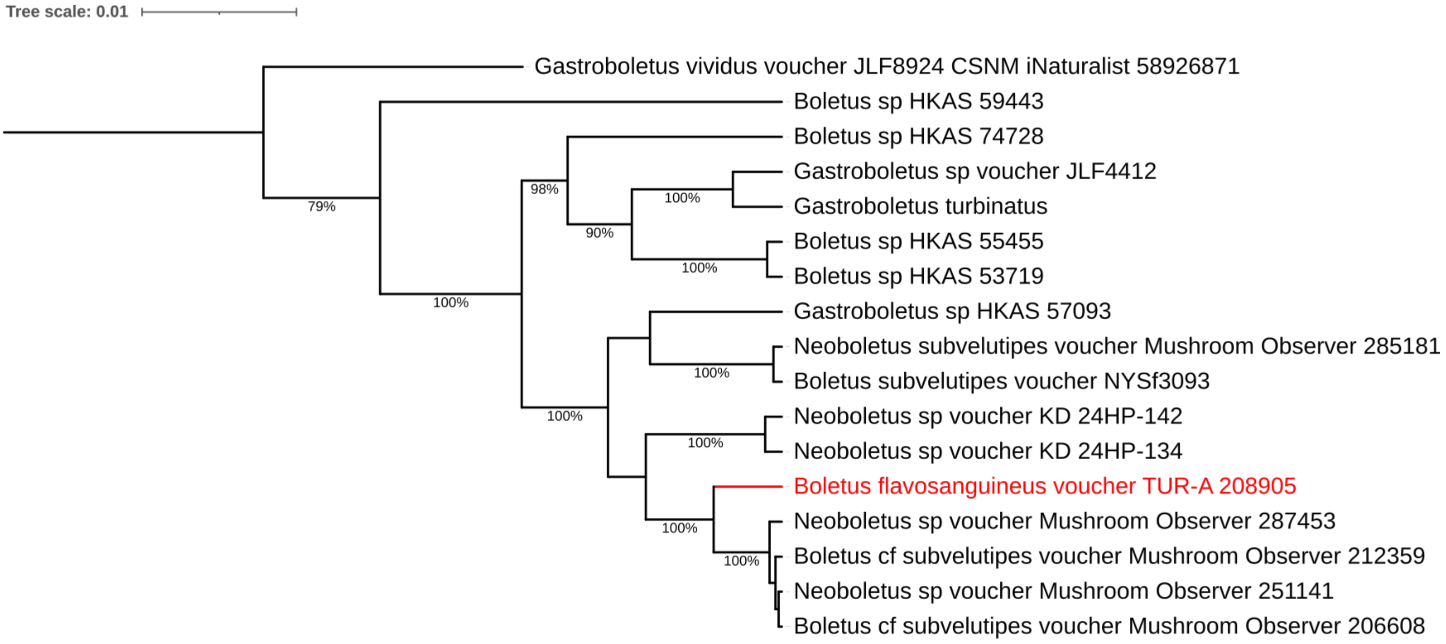

### Supplementary Figures Legends

**Fig. S1** (Online Resource 2) Phylogeny for representatives of the subfamily Suillelloideae, based on a concatenate of nucleotide alignments of the nucLSU, RPB1, RPB2 and TEF1 genes. Branches are coloured according to the preceding taxonomy: *Cyanoboletus* s.l. (dark blue), *Acyanoboletus* (light blue), *Cupreoboletus* (purple) and *Cacaoporus* (brown). The position of *Neoboletus flavosanguineus* is highlighted in red. The tree prior is constant size coalescent, relaxed clock, and $2.5 \times 10^7$ states were obtained, with a burnin of $5 \times 10^6$ states. Joint ESS 26.3. MrHIPSTR tree's log clade credibility: -17.2327. Number of clades with credibility > 0.5: 61.

**Fig. S2** (Online Resource 3) Single-gene phylogenies for a) nucLSU b) RPB1 c) RPB2 and d) TEF1. Branches colorised according to the genus assignment in Fig. 1: dark blue, *Cyanoboletus*; light blue, *Cacaoporus*; purple, *Cupreoboletus*. In all cases a strict clock was used. A total $1.5 \times 10^7$ states were obtained in a), with a burnin of $10^6$ states. Joint ESS: a) 1019, b) 2673.5, c) 447.6, d) 852. MrHIPSTR tree's log clade credibility: a) -90.2592, b) -16.4816, c) -46.3084, d) -40.4888. Number of clades with credibility > 0.5: a) 52, b) 24, c) 57, d) 56.

**Fig. S3** (Online Resource 4) Phylogeny for a thorough sampling of nucLSU sequences concatenated with matching ITS sequences, taken from the alignments pertaining to Online Resource 3a and Fig. 1, respectively. Colour codes as in Fig. 1 and Online Resource 3. Putative clade epithets at species level not available in Fig. 1 are shown. Asterisk indicates the '*pulverulentus sensu lato*' clade. A relaxed clock was used, and a total of $3.6 \times 10^7$ states were obtained after combining those from each run, with a burnin of $2 \times 10^6$ states each. Joint ESS 144.9. MrHIPSTR tree's log clade credibility: -68.4869. Number of clades with credibility > 0.5: 61.

**Fig. S4** (Online Resource 5) Phylogeny for a thorough sampling of RPB2 and TEF1 sequences concatenated with matching nucLSU sequences, taken from the alignments pertaining to Online Resource 3c/d/a, respectively. Colour codes as in Fig. 1 and Online Resource 3. Specimen additions to Fig. 1 and Online Resource 3 are indicated. A relaxed clock was used, and a total $10^7$ states were obtained, with a burnin of $2.2 \times 10^6$ states. Joint ESS 43.4. MrHIPSTR tree's log clade credibility: -37.6367. Number of clades with credibility > 0.5: 66.

**Fig. S5** (Online Resource 6) Phylogeny for the ITS based on pairwise comparisons, representative of collapsing clades at gap threshold 0.6%. Accession numbers KY548897, KY548911, MH257571, and MT345198 used in Fig. 1 are not included here, being too short for meaningful divergence calculations. Branch supports obtained from 5000 bootstraps. For clarity, bootstrap support values below 60% are not shown.

**Fig S6** (Online Resource 8) Approximate geographic distributions of the sequenced collections of *Cyanoboletus pulverulentus* (violet dots) and *Cy. mediterraneensis* (red dots). Built in https://www.mapcustomizer.com (Ursus Software, LLC).

**Fig. S7** (Online Resource 9) Phylogenies around *Neoboletus flavosanguineus* a) ITS (strict clock), b) concatenate of nucLSU, RPB2 and TEF1 alignments (relaxed clock). MrHIPSTR tree's log clade credibility: a) -25.6862 b) -4.6686. Number of clades with credibility > 0.5: a) 18 b) 12
